# Human CCR6^+^ Th cells show both an extended stable gradient of Th17 activity and imprinted plasticity

**DOI:** 10.1101/2023.01.05.522630

**Authors:** Satya P. Singh, Farhat Parween, Nithin Edara, Hongwei H. Zhang, Jinguo Chen, Francisco Otaizo-Carrasquero, Debby Cheng, Nicole A. Oppenheim, Amy Ransier, Wenjun Zhu, Amirhossein Shamsaddini, Paul J. Gardina, Samuel W. Darko, Tej Pratap Singh, Daniel C. Douek, Timothy G. Myers, Joshua M. Farber

## Abstract

Th17 cells have been investigated in mice primarily for their contributions to autoimmune diseases. However, the pathways of differentiation of Th17 and related (type 17) cells and the structure of the type 17 memory population in humans are not well understood; such understanding is critical for manipulating these cells *in vivo*. By exploiting differences in levels of surface CCR6, we found that human type 17 memory cells, including individual T cell clonotypes, form an elongated continuum of type 17 character along which cells can be driven by increasing RORγt. This continuum includes cells preserved within the memory pool with potentials that reflect the early preferential activation of multiple over single lineages. The CCR6^+^ cells’ phenotypes and epigenomes are stable across cell divisions under homeostatic conditions. Nonetheless, activation in polarizing and non-polarizing conditions can yield additional functionalities, revealing, respectively, both environmentally induced and imprinted mechanisms that contribute differentially across the continuum to yield the unusual plasticity ascribed to type 17 cells.

## INTRODUCTION

Immunological memory mediated by the adaptive immune system can provide long-standing natural and vaccine-mediated protection against infectious agents. Notwithstanding controversies regarding the meaning and effectiveness of the T cell component of immunological memory (1), pathogen-specific CD8^+^ and CD4^+^ T cells can be detected in humans many decades following exposure (2, 3), and for CD4^+^ memory T cells, which are the focus of the current studies, there are data from mice demonstrating their protective capabilities (4, 5). T cell memory depends on epigenetic memory, including DNA methylation, with resting memory T cells maintaining an altered epigenome that is poised to allow cells to respond to activation in a biased but flexible fashion based on the pattern of initial differentiation/polarization (6).

The ontogeny of memory T cells has long been a subject of ongoing investigation and debate (7). It is clear that descendants of individual CD8^+^ T cells can assume multiple phenotypes of effector and memory cells (8-10). Further, based on studies of human CD4^+^ memory T cells specific for pathogens or tetanus toxoid, individual T cells can, and often do give rise to cells of different lineages as defined by Th1, Th2, and Th17 signature cytokines (11). Moreover, the preponderance of evidence from mouse and human studies favors effector T cells as the direct precursors of memory cells (8, 12-18). Critical experiments using single-cell transfers in mice have shown that the ability to form memory cells is a general and not exceptional property of individual naive CD4^+^ T cells, and that, of equal importance, the composition of the memory population mirrors the composition of the effector population (18). Moreover, studies of human memory CD8^+^ T cells have suggested that phenotypes and epigenomes are stable under homeostatic conditions (19). The correspondence between effector and memory populations provides a fundamental benefit often lacking in models explaining the origins of T cell memory, namely that the information contained in the acute and evolving effector response is fully preserved in the memory population. A further consequence of these findings is that the memory T cell population preserves a record of the acute response and can be used to reconstruct pathways of differentiation taken by activated T cells. The ability to use studies of the memory population in this way is particularly important in understanding the process of memory T cell differentiation in humans, where it is both the case that the time frames and complexity of environmental exposures far exceed those of laboratory mice (20, 21), and detailed observations of immune responses in real time are difficult.

Based on the conventional definitions of CD4^+^ T cell effector/memory subsets, Th17 cells were the third major subset to be identified as a “lineage”, with IL-17A as their signature cytokine and RORγt as their signature transcription factor (22). In initial studies in mice, Th17 cells were found to contribute to autoimmune pathology (23), and subsequent work in humans demonstrated the importance of the Th17 pathway in host defense against *Candida, Staphylococcus aureus*, and *Mycobacterium* (24-27) as well as in inflammatory disease (28). Further, mouse studies revealed that Th17 cells can assume dichotomous phenotypes as both protective and pathogenic (29) and that although some studies in mice have questioned the durability of Th17 memory (30), other studies in mice (31) and humans (32) have reported that Th17 cells can establish persistent memory.

Prominent features described for mouse Th17 cells include their abilities to co-express genes of alternative lineages, lose expression of IL-17A/F, and acquire the ability to express a different signature cytokine, primarily IFNγ (33-37) - manifesting properties broadly described as plasticity (37). Nonetheless, whether plasticity in cytokine profiles indicates that a fully differentiated memory cell is able to transdifferentiate remains controversial (38), and some studies of Th17 cells in mice have shown that their phenotype can be stable during both homeostasis and inflammation (39, 40). Investigations of the phenotype and epigenome of human Th17 cells suggest that they may be more stable than their murine counterparts (41), and that IFNγ^+^IL-17A^-^ cells that arise under Th17 conditions are best viewed as a subset of Th17 cells rather than as Th1 cells that have arisen by transdifferentiation (42, 43).

The chemokine receptor CCR6 and its ligand, CCL20, have broad roles in humoral (44) and cellular (45) barrier defense. We and others reported that virtually all human Th17 cells express CCR6 (46-48), which is inducible on T cells by RORγt (49). Cells expressing other cytokines whose expression is associated with, but not precisely matching the expression of IL-17A, such as IL-22 and CCL20, are also found exclusively as subsets of CCR6^+^ cells (46, 50) and this manuscript); and CCR6 is found on a subset of human peripheral regulatory T cells (51) that can be stimulated to produce IL-17A (52). Additionally, CCR6 is used to identify the human Th17-related, IFNγ^+^IL-17A^-^ cells that have been referred to as nonclassical Th1 (43) or Th1* (46) cells, and that have been implicated in defense against mycobacteria (26) and autoimmune disease (53). Together, these data suggest that human Th17 cells that are defined by their production of IL-17A/F represent a limited subset within a broader family of “type 17”, CCR6-expressing cells. Our studies started from the premise that investigating the interrelationships of cells within this larger type 17 family would reveal how human Th17 and Th17-related effector/memory cells are formed and organized, thereby providing additional insights into their roles in host defense and inflammation.

We discovered previously in analyzing relationships between the expression of signature chemokine receptors and a limited number of other lineage-associated genes in bulk populations of human Th cells that levels of surface CCR6 correlated positively with levels of expression of *IL17A* and *RORC*, and that the CCR6^+^ cells showed a wide gradient in their expression of these genes as compared with gradients for expression of type 1 genes in CXCR3^+^ cells or type 2 genes in CCR4^+^ cells (54). In the current study, by analyzing human CD4^+^ T cells based on their levels of surface CCR6, both as populations and as single cells, we have found that the CCR6^+^ cells form an extended, progressive continuum of type 17 character, with cells of individual T cell clonotypes found throughout the continuum, albeit with changing frequencies. Although cells could be driven along the continuum by over-expression of *RORC*/RORγt, analysis of genome-wide patterns of CpG methylation in dividing cells showed that epigenotypes/phenotypes are stable under homeostatic conditions. Single-cell analysis suggested pan-potentiation of effector pathways early after activation of CD4^+^ T cells *in vivo* that persist within the memory population, and most type 17 cells can co-express genes of alternative lineages, with co-expression diminishing along the type 17 continuum. Restimulation of CCR6-defined subgroups under polarizing and non-polarizing conditions, along with over-expression of lineage-defining transcription factors, could yield additional functionalities by both environmentally induced and previously imprinted mechanisms that contributed differentially across the continuum to type 17 cell plasticity.

## Materials and methods

### Human samples

Elutriated lymphocytes and whole blood were obtained from healthy donors by the Department of Transfusion Medicine, National Institutes of Health, under an Institutional Review Board-approved protocol.

### Purification of lymphocyte subsets from blood and cell sorting

CD4^+^ T cells were isolated from elutriated lymphocytes by negative selection using RosetteSep CD4^+^ T cell enrichment cocktail (StemCell Technologies) followed by Ficoll/Hypaque (Amersham Biosciences) density centrifugation according to the protocol from StemCell Technologies. If necessary, red blood cells were removed using Pharm Lyse (BD Biosciences) according to the manufacturer’s protocol. For staining cells were suspended in 250 μl of Hanks Balanced Salt Solution (HBSS, Mediatech) containing 4% fetal bovine serum (FBS, GeminiBio) and incubated with following antibodies: anti-CD4, anti-CD45RO, and anti-CCR6 for 30 min at room temperature. In some experiments, in addition to these antibodies, cells were incubated with anti-CD25 and anti-CD62L, anti-CXCR5, anti-CXCR3 and anti-CCR4 antibodies. All antibodies were against human antigens. Cells were washed and resuspended in HBSS plus 4% FBS. CD4^+^ T cells were sorted into CD45RO^−^ (naïve), and CD45RO^+^ subsets, and the CD45RO^+^ subset was further separated into CCR6^neg^, CCR6^pos^, and, within the CCR6^pos^ cells, into CCR6^low^, CCR6^int^ and CCR6^high^ subsets. For *in vitro* polarization experiments, cells were sorted into naïve (CD4^+^CD45RO^−^CD25^−^CXCR5^−^CCR4^−^ CXCR3^−^), CCR6^neg^ (CD4^+^CD45RO^+^CD25^−^CXCR5^−^CXCR3^−^CCR6^−^ or CD4^+^CD45RO^+^CD25^−^ CXCR5^−^ CCR4^−^CCR6^−^ or CD4^+^CD45RO^+^CD25^−^CXCR5^−^CXCR3^−^CCR4^−^CCR6^−^), CCR6^low^ (CD4^+^CD45RO^+^CD25^−^CXCR5^−^CXCR3^−^CCR6^low^ or CD4^+^CD45RO^+^CD25^−^CXCR5^−^CCR4^−^ CCR6^low^ or CD4^+^CD45RO^+^CD25^−^CXCR5^−^CXCR3^−^CCR4^−^CCR6^low^), and CCR6^high^ (CD4^+^CD45RO^+^CD25^−^CXCR5^−^CXCR3^−^CCR6^high^ or CD4^+^CD45RO^+^CD25^−^CXCR5^−^CCR4^−^CCR6^high^ or CD4^+^CD45RO^+^CD25^−^CXCR5^−^CXCR3^−^CCR4^−^CCR6^high^). All cell sorting was done using a FACSAria Cell Sorter (BD Biosciences). By post-sort analysis, the purity of populations was 95-99%. Cells were used for experiments immediately after sorting.

### Isolation of RNA, and real-time fluorogenic RT-PCR

T cells were activated with 20 ng/ml PMA and 1 mM ionomycin in RPMI 1640 (GIBCO-Thermo Fisher Scientific) supplemented with 10% FBS, 1% L-glutamine and 1% penicillin/streptomycin (complete medium) for 3 h at 37°C under 5% CO_2_ and total cellular RNA was purified using PureLink™ RNA kit (Life Technology) or RNeasy Mini Kit (Qiagen) and the concentration and RNA Integrity Number (RIN) was determined with a RNA 6000 nano kit on 2100 Bioanalyzer instrument (Agilent Technologies). Real-time PCR was performed on samples in duplicate on an ABI 7900HT Real-Time PCR System (Applied Biosystems) or Eco Real-Time PCR System (Illumina) using SuperScript One Step RT-PCR kit (Life Technology). For most experiments, primer and probe sets (FAM/MGB-labeled) were purchased from Applied Biosystems. In some cases, DELTAgene Assay primers were purchased from Fluidigm and amplicon DNA was quantified using SsoFast™ EvaGreen Supermix with Low ROX (Bio-Rad Laboratories). Gene expression data were normalized based on the values for *GAPDH* mRNA. For some assays, for cells from each donor, relative levels of expression, based on values for 2^−ΔCT^, are shown after normalization to the single highest value, which was set to 100. For other assays, values of 2^−ΔCT^ or ΔCT are shown without additional normalization or shown as fold changes relative to the values for naïve cells, which were set at 1.

### Intracellular and intranuclear staining for flow cytometry

T cells purified by cell sorting were stimulated with Leukocyte Activation Cocktail, with GolgiPlus™ (BD Pharmingen) for 6 h at 37°C under 5% CO_2_ before being stained with following antibodies alone or in combinations: anti-IL-17A (eBioscience), anti-IL-17F, anti-IL-22, anti-IL-6, anti-CCL20 and anti-IFNγ (R&D Systems) by using Cytofix/CytoPerm Plus kit (BD Pharmingen) or anti-RORγt, anti-FOXP3, anti-T-bet or anti-GATA3 using Foxp3 staining buffer set (eBioscience) according to the manufacturer’s instructions. All flow cytometry was done using FACSCaliber, LSR II System or Fortessa flow cytometers (BD Biosciences), and the data were subsequently analyzed using FlowJO software (TreeStar).

### ELISA (Enzyme-Linked Immunosorbent Assay) for cytokine and chemokine production

T cells purified by cell sorting were activated with 20 ng/ml PMA and 1 mM ionomycin in complete medium at 37°C under 5% CO_2_ for 12 h and supernatants were collected and stored in -80°C freezer until analyzed. GM-CSF, TNF-α, IFNγ, IL-4, IL-5, IL-10, IL-13, IL-17A, CCL3, CCL4, CCL5 and CCL20 were measured using Search Light Proteome Array (Pierce Biotechnology).

### Microarray analyses

FACS-sorted naïve, CCR6^neg^, CCR6^low^, CCR6^int^ and CCR6^high^ subsets of CD4^+^ T cells were used for isolating RNA without and with prior activation with 20 ng/ml PMA and 1 mM ionomycin for 3 h at 37°C under 5% CO_2_ using RNeasy Mini Kit (Qiagen) and gene expression levels were determined using Illumina Human WG6 Expression BeadChip Version 3 arrays (Illumina Inc.) according to the manufacturer’s protocol. In brief, 500 ng of total RNA was used to make cDNA and label cRNA using the Ambion Illumina TotalPrep RNA Amplification kit (Invitrogen). The resulting cRNA samples were hybridized to an Illumina Human WG6 Expression BeadChip array for 14-20 hour at 58°C in an Illumina Hybridization Oven (Illumina Inc.). Arrays were washed and scanned following the protocols in the Illumina Whole-Genome Gene Expression Direct Hybridization Assay Guide (Illumina). Signal data were extracted from the image files with the Gene Expression module of the GenomeStudio software (Illumina, Inc.) Signal intensities were converted to log_2_ scale. Detection *p* values were calculated as described in the GenomeStudio Gene Expression Module User Guide. Data for array probes with insufficient signal were removed from the dataset (detection was defined as at least two arrays with detection *p* values < 0.1). Analysis of variance (ANOVA) was then performed on the normalized log_2_ expression estimates to test for mRNA expression differences across cell subgroups, with contrasts for the CCR6^low^, CCR6^int^ and CCR6^high^ subsets verses the CCR6^neg^ subgroup. A *p* value of 0.05 was used for the significance cutoff, after adjusting for the familywise error rate (FWER) using the Benjamini-Hochberg method to account for multiple testing. One-way hierarchical clustering (on probes, i.e., gene-wise) was performed on the standardized LS means for all genes whose differential expression reached significance, separately for stimulated and non-stimulated cells. Genes whose differential expression reached significance and whose differences were the most extreme (greater than 4 standard deviations (|Z| > 4) of log_2_ differences within any contrast) were then analyzed to determine patterns across the CCR6-expressing subsets. Log_2_ differences from the contrasts were standardized across each probe and the resulting Z-scores were clustered by K-means where k=12. Genes within each cluster for each contrast were summarized as mean Z-scores and shown as a heat map to illustrate relative expression patterns across the CCR6-expressing subsets. Statistical analyses were performed using JMP/Genomics software version 7.0 (SAS Institute Inc.).

### Genome-wide DNA methylation analysis

Genomic DNAs from FACS-sorted naïve, CCR6^neg^, CCR6^low^, CCR6^int^ and CCR6^high^ subgroups were purified using the DNeasy kit (Qiagen). In some experiments, after purification by cell sorting, a portion of cells were used as starting cells (day 0) and the remaining cells were labeled with CFSE and cultured with the homeostatic cytokines IL-7 and IL-15, for 7 days. Cells from the different subgroups were again sorted, based on CFSE dilution, into non-proliferated and proliferated cells and genomic DNAs were purified using the DNeasy kit. Bisulfite conversion of genomic DNA (500 ng) was performed using the Zymo Research EZ DNA Methylation kit (Zymo Research) according to the manufacturer’s recommendations for the Illumina Infinium assay. The bisulfite conversion incubation was processed using alternative thermocycler settings as outlined in the “Alternative Incubation Conditions When Using the Illumina Infinium® Methylation” section of the Zymo instruction manual appendix. The bisulfite-converted DNA samples were then processed for hybridization and staining using either the Illumina Infinium Human Methylation 450 BeadChip Kit or the Illumina Infinium MethylationEPIC BeadChip Kit (Illumina Inc.) according to the manufacturer’s recommendations. DNA methylation ratios were calculated on a scale of 0 - 1 in which zero represent 100% unmethylated and 1 represent 100% methylated. The methylation dataset was visualized using Tidyverse (an R open-source package).

### Single-cell mRNA sequencing

Purified CD4^+^ T cells were FACS sorted into CCR6^neg^, CCR6^low^, CCR6^int^ and CCR6^high^ subgroups, loaded with CFSE, and cultured with autologous dodecyldimethylamine oxide (DDAO)-loaded, irradiated PBMCs in the presence of an extract of CMV-infected cells (EastCoast Bio, North Berwick, ME). On day 7, the proliferated, CFSE^low^ T cells were sorted, counted for cell viability, and immediately used for library preparation using Chromium™ Single Cell V (D) J Reagent Kit (10X Genomics) according to the manufacturer’s instructions.

The single-cell TCR libraries were sequenced on an Illumina NextSeq500 to a minimum sequencing depth of 5000 reads per cell using paired-end sequencing with single indexing as follows: read length of 150 bp read 1, 8 bp i7 index, 150 bp read 2. The raw data were downloaded using Python Run Downloader (Illumina) and analyzed using Cellranger (Illumina). TCR repertoires were processed using custom scripts to produce Circos-compatible files. In each Circos plot (Circos, version 0.69), the relative size of a clonal expansion is represented by arc-length while overlapping clonotypes are represented by ribbons connecting arcs. In some cases, to increase interpretability of the Circos plots, ribbons were only drawn between clonotypes that represented 4% or greater of either of the connected TCR repertoires. Morisita-Horn Dissimilarity values between TCR repertoires were calculated using the R package vegan (v. 2.5-7) and plotted in R using reciprocal values to represent similarity.

The single-cell 5’ gene expression libraries were sequenced on Illumina NextSeq500 to a minimum sequencing depth of 50000 reads per cell using pair-end sequencing as follows: read length of 150 bp read 1, 8 bp i7 index, 150 bp read 2. The raw data were downloaded using Python Run Downloader (Illumina) and analyzed using Cell Ranger (Illumina).

### *TCRB* deep sequencing

CD4^+^ T cells were sorted into naive, CCR6^neg^, CCR6^low^, CCR6^int^ and CCR6^high^ subgroups and total cellular RNAs were purified from using PureLink™ RNA kit (Life Technology) and the concentrations and RNA Integrity Numbers (RIN) were determined with an RNA 6000 nano kit on a 2100 Bioanalyzer (Agilent Technologies). Libraries for TCR sequencing were prepared as described (55). TCRB annotation was performed by first determining which paired reads were of the expected library insert size by merging the paired-end reads with the bioinformatic tool, PEAR (56). Paired-end reads that could be merged were discarded as they represented library inserts that were too short and assumed to be off-target products. Unmerged read-pairs were then annotated with MiXCR (57). Heatmaps and Circos plots were then generated in the same manner as above.

### High-throughput single cell gene expression analysis by microfluidic quantitative RT-PCR

Individual cells from naïve, CCR6^neg^, CCR6^low^, CCR6^int^ and CCR6^high^ subgroups were sorted by FACS into single wells of a 96 well plate where each well contained 5 ul of CellDirect 2X reaction mixture solution (Invitrogen) supplemented with 0.05 units of SUPERase-In™ RNase inhibitor (Ambion). After incubating at 4°C for 15 min, each plate containing sorted cells was frozen and stored at -30° C until analysis. For analysis, plates were thawed on ice and each well was supplemented with 0.2 ul SuperScript III RT Platinum Taq Mix (Invitrogen), 2.8 ul DNA suspension buffer (Invitrogen) and 1.0 ul of a mixture of 48 pooled DELTAgene Assay primer pairs containing each primer at a final concentration of 500 nM (Fluidigm). mRNAs from lysates of single cells were reverse transcribed into cDNA at 50° C for 15 min, followed by inactivation of the reverse transcriptase at 95° C for 2 min. cDNAs were pre-amplified for 20 cycles by denaturing at 95° C for 15 sec followed by annealing and amplification at 60° C for 4 min. To digest unincorporated primers, 3.6 ul of Exonuclease I reaction mixture containing 0.36 ul of 10X Exonuclease reaction buffer, 0.72 ul of 20 units/ul Exonuclease I (New England BioLabs) and 2.52 ul of nuclease free water was added to each well, followed by an incubation at 37° C for 30 min and finally treatment at 80° C for 15 min. The pre-amplified cDNA was diluted 5-fold with TE buffer (10 mM Tris, pH 8.0, 1 mM EDTA) prior to gene expression assays using the 96.96 (M96) Dynamic Array Microfluidic Chip on a BioMark HD System (Fluidigm) according to the manufacturer’s protocol. Briefly, 2.25 ul of pre-amplified cDNA was mixed with 2.5 ul of 2X SsoFast EvaGreen Supermix with low ROX (Bio-Rad) and 0.25 ul of 20X DNA binding dye sample loading reagent (Fluidigm) and transferred into one of the array microfluidic chip sample inlets. Individual DELTAgene assays were prepared by combining 0.25 ul of primer pairs (100 uM), 2.25 ul of DNA suspension buffer, 2.5 ul of 2X Assay loading reagent (Fluidigm) and transferred into one of the array microfluidic chip assay inlets. Samples and assay mix solutions were loaded on the microfluidic chip using an IFC Controller MX (for 48.48 Dynamic array IFC) or IFC Controller HX (for M96 Dynamic array IFC) (Fludigm) and transferred to a BioMark HD System and run following manufacturer’s protocol. For each subgroup, cells were sorted from three individual donors. All Ct values obtained from the BioMark HD System were normalized to the endogenous control genes, *GAPDH* and *PGK1* and cells with very low (Ct > 30) or non-detected endogenous control genes were removed from the analysis. For further analysis, normalized Ct values for individual genes were calculated by subtracting from the Ct value the mean Ct values of that gene on all the samples from a given subgroup on an individual chip and dividing by 3 times the standard deviation as described (58). Data acquired from the BioMark HD system were processed using Singular Analysis Toolset Software v3.5.2 (Fluidigm). We used open-source R packages (Tidyverse) to process the datasets and visualization. In some experiments, pre-amplified samples were used for TaqMan quantitative PCR which was performed on Eco Real-Time PCR System (Illumina) using gene specific primers purchased from Fluidigm (DELTAgene Assay) and detected using SsoFast™ EvaGreen Supermix with Low ROX (Bio-Rad Laboratories). Results from these assays were normalized based on the values for *GAPDH* mRNA.

### Ectopic *RORC, TBX21* and *GATA3* expression by lentiviral gene transduction

All the lentivirus particles expressing sequences from *RORC, TBX21* and *GATA3* or control virus particles contained the pReceiver-LV201 vector and were obtained from GeneCopoeia. A CMV promoter was used to drive the *RORC, TBX21* and *GATA3* sequences and an SV40 promoter was used to drive eGFP sequences. Purified CD4^+^ T cells were sorted by FACS into memory cells that were CCR6^neg^, CCR6^low^, and CCR6^high^ and transduced with lentivirus particles at an MOI of 5 using retronectin (TaKaRa) according to the manufacturer’s instructions. In brief, 1 × 10^6^ sorted cells per well of a 24 well plate were activated with anti-CD3/anti-CD28-coated beads, 1 bead per cell, for 24 hours and transferred to a well precoated with retronectin (TaKaRa). Lentivirus transduction was performed by spinoculation at 1200 X g for 2 hours at 32° C in the presence of virus particles and 8 ug/ml polybrene (Millipore). The next day, virus particles were removed, and cells were cultured in complete RPMI-1640 supplemented with 10 ng/ml of IL-2 for an additional three days. Cells were stimulated with PMA and ionomycin for 3 h and GFP^+^ cells were sorted by FACS. For single cell gene expression experiments, cells were sorted by FACS into each well of a 96 well plate containing 5 ul of CellDirect 2X reaction mixture solution supplemented with 0.05 units of SUPERase-In™ RNase inhibitor and used for analysis using the M96 Dynamic Array Microfluidic Chip on a BioMark HD System according to the manufacturer’s protocol or analyzed using quantitative RT-PCR. For some experiment cells were loaded with DDAO before lentivirus transduction to enable cells to be sorted based on numbers of cell divisions.

### *In vitro* polarization of CD4^+^ T lymphocytes from adult peripheral blood

Cells were sorted by FACS as described above into naïve and CCR6^neg^, CCR6^low^ and CCR6^high^ memory subgroups and cultured at 1 × 10^6^ cells/ml in 24 well plates in ImmunoCult XF T cell expansion medium (Stem Cell Technologies) supplemented with 10% FBS and IL-2 (200 units/ml). Stimulation was done using ImmunoCult human CD3/CD28/CD2 T Cell Activator (Stem Cell Technologies) in non-polarizing conditions using anti-IL-4 (5 *μ*g/ml), and anti-IFN-γ (10 *μ*g/ml), or in Th1 conditions using recombinant human IL-12 (5 ng/ml), and anti-IL-4 (4 *μ*g/ml), or in Th2 conditions using recombinant human IL-4 (10 ng/ml), and anti-IFN-γ (10 *μ*g/ml). After three days, proliferating cells were washed in PBS and expanded under the same conditions in the absence of CD3/CD28/CD2 T Cell Activator.

### Quantification and statistical analysis

Data were analyzed using GraphPad Prism software version 9 (GraphPad, San Diego, CA). The *p* values were calculated using Wilcoxon matched pairs signed rank test. Linear regression in R was used to generate p-values for the relationship between single-cell q-PCR Ct values for genes of interest. ANOVA was performed on the normalized log_2_ expression values to estimate differences between cell subgroups, and adjusted *p*-values were generated using the Benjamini-Hochberg method to account for multiple testing. \**p* < 0.05, \*\**p* < 0.01, and \*\*\*\**p* < 0.0001.

## RESULTS

### CCR6-defined subgroups show progressive upregulation of type 17, and down-regulation of type 1 and type 2 genes and proteins

To investigate the relationship between the surface expression of CCR6 and genes associated with effector/memory phenotypes, we used FACS to separate CD4^+^ T cells from healthy adult blood donors into naïve cells and memory-phenotype cells that were CCR6^neg^ and CCR6^pos^, and the CCR6^pos^ cells were further separated into three equal subgroups based on their levels of CCR6: CCR6^low^, CCR6^int^ or CCR6^high^ (Fig. 1 A). Intracellular staining of FACS-purified cells activated with PMA and ionomycin revealed that virtually all cells expressing type 17 cytokines/chemokines, such as IL-17A, IL-17F, IL-22 and CCL20 are found within CCR6-expressing subset (Fig. S1, A and B). Building on our previous studies (54), using data from gene arrays in cells from eight donors, we analyzed changes in gene expression in cells that were either left non-stimulated or stimulated *ex vivo* with PMA and ionomycin. As shown in volcano plots in Figure 1B, in comparing the CCR6^+^ subgroups to CCR6^neg^ cells as a common comparator, we found that type 17 genes, such as *IL17A, IL17F, IL22, IL23R, CCL20, RORA* and *RORC* (shown in red), were progressively upregulated, and type 1 and type 2 genes, such as *IFNG, CXCR3, CCL3, CCL5, IL4* and *IL5* (shown in green) were progressively downregulated in CCR6^low^ to CCR6^int^ to CCR6^high^ cells. Pairwise comparisons of gene-specific probes among the subgroups revealed approximately 600 genes that were differentially expressed in any of the comparisons for the non-stimulated and/or stimulated cells based on a false discovery rate of 0.05. It is important to note, as can be appreciated from the mRNAs of diminishing abundance across the subgroups, that the CCR6^neg^ subgroup is particularly heterogeneous, containing, for example highly differentiated Th1 and Th2 cells. All differentially expressed genes were grouped by k-means into twelve clusters with a variety of patterns across the CCR6^low^, CCR6^int^ and CCR6^high^ subgroups. Genes in most clusters display monotonic changes in expression. A salient gene within each cluster is shown, such as the signature genes for the Th lineages, *IL17F, IFNG*, and *IL5* (Fig. 1 C). Heat maps of gene expression from the naïve and CCR6-defined (CCR6^neg^, CCR6^low^, CCR6^int^ and CCR6^high^) memory subgroups, both non-stimulated and stimulated, show large differences in patterns between naïve and memory cells and between CCR6^neg^ and CCR6^pos^ cells and progressive changes from CCR6^low^ to CCR6^high^ cells (Fig. S1 C).

**Figure 1.**
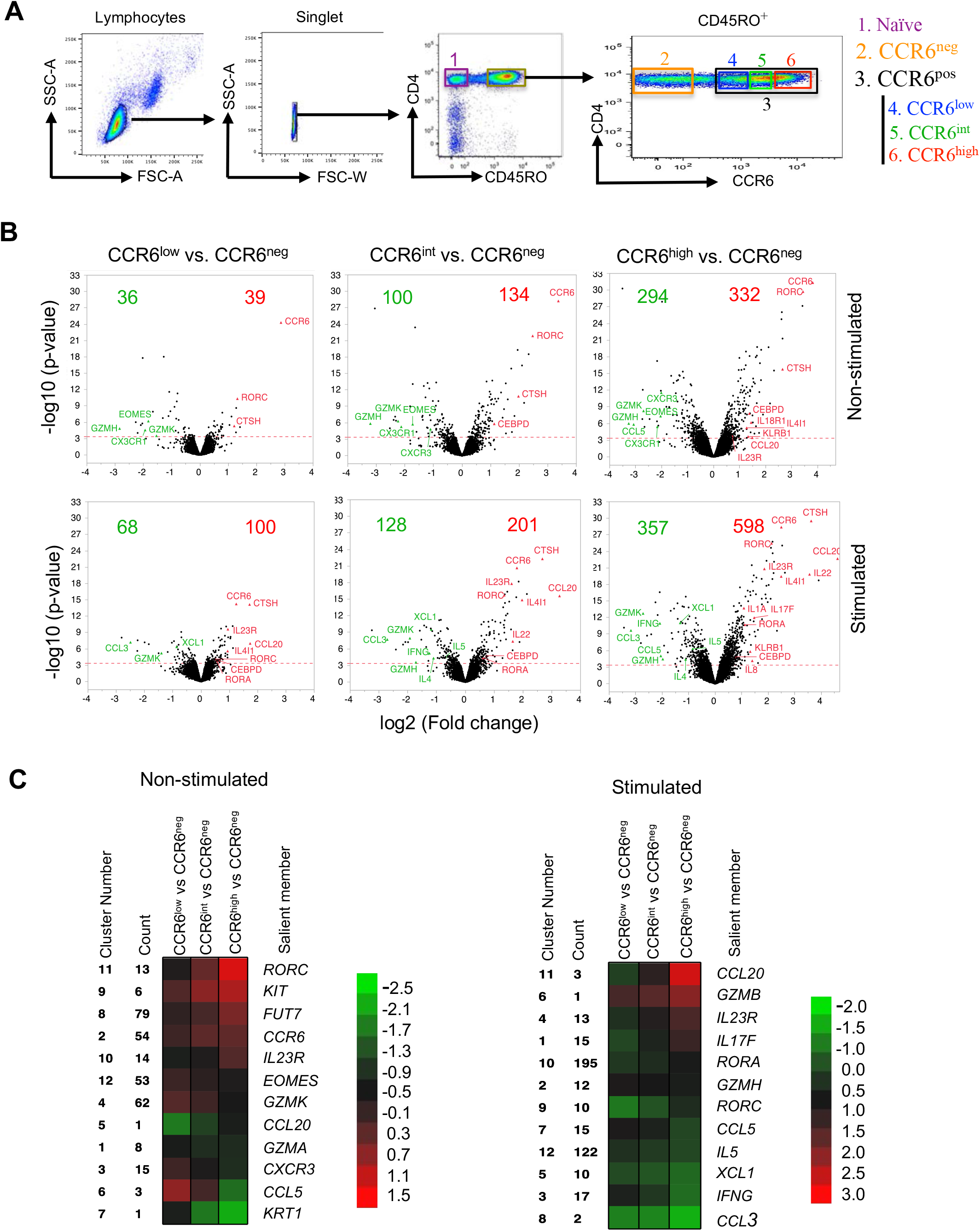
Progressive upregulation of type 17 genes and downregulation of type 1 and type 2 genes in CCR6-expressing memory subgroups. **(A)** Elutriated lymphocytes from blood that had been enriched for CD4^+^ T cells were separated based on CD45RO staining into CD45RO^−^ (naïve) and CD45RO^+^ (memory) cells and the CD45RO^+^ cells were either separated into CCR6^neg^ and CCR6^pos^ subgroups or into CCR6^neg^, CCR6^low^, CCR6^int^ and CCR6^high^ subgroups. **(B)** Volcano plots showing fold changes (log_2_ ratios) versus significance (-log_10_ *p*-values) for genes in non-stimulated (top) or stimulated (bottom) cells for CCR6^low^, CCR6^int^ and CCR6^high^ subgroups compared with the CCR6^neg^ subgroup. The red dotted line marks the cutoff for significance of differential expression at a false discovery rate ≤0.05. On each volcano plot are indicated the number of genes upregulated (red) or downregulated (green), and some genes of interest are labeled. **(C)** Heatmaps illustrating the relative patterns of gene expression differences between CCR6^low^, CCR6^int^ and CCR6^high^ subgroups compared with the CCR6^neg^ subgroup. The standardized differences for the gene-specific probes showing the greatest differences in these comparisons were clustered by K-means (Figure S2); the color/value in each small box represents the mean across the genes in the cluster for each contrast. The cluster number, the number of unique gene-specific probes in that cluster and a representative gene in each cluster are shown. Analyses shown are combined from eight individual donors.

We next performed a more targeted analysis of the gradients of lineage-related genes and proteins across the naïve and CCR6-defined memory subgroups. Consistent with the gene array data, real-time RT-PCR of cells activated with PMA and ionomycin showed increases in expression of type 17 genes and decreases in signature type 1 (*IFNG* and *TBX21*) and type 2 genes (*IL4, IL5, IL13* and *GATA3*) across the CCR6-defined memory subgroups (Fig. 2 A). Similar patterns were found for lineage-associated proteins as detected by intracellular staining or ELISA (Fig. 2 B and Fig. S2 A). We further investigated pair-wise relationships among type 17 proteins across the naïve and CCR6-defined memory subgroups by intracellular staining. For IL-17F, as depicted in Figures S1 A and S2 B, virtually all positive-staining cells were found as a subset of cells expressing IL-17A. For IL-17A/F vs. CCL20, both CCL20^+^IL-17A/F^−^ cells and CCL20^+^IL-17A/F^+^ cells increased from the CCR6^low^ to the CCR6^high^ subgroup, although the fold increase in CCL20^+^IL-17A/F^+^ cells was greater across the subgroups, leading to an increase in the ratio of IL-17A/F^+^CCL20^+^: IL-17A/F^−^CCL20^+^ cells across the subgroups. For IL-17A vs. IL-22 there was a significant increase in the frequency of IL-22^+^IL-17A^+^ cells from the CCR6^low^ to the CCR6^high^ subgroup (although the ratio of IL-17A^+^ IL-22^+^: IL-17A^+^ IL-22^−^ cells was constant across the subgroups), but very little change in the frequency of IL-22^+^IL-17A^−^ cells suggesting that IL-22 may be upregulated through different pathways in cells that do vs. cells that do not co-express IL-17A. For CCL20 vs. IL-22, based on the percentages of single-positive vs. double-positive cells, their increases across the subgroups showed no evidence of being co-regulated (Fig. S2 B). In contrast to the patterns for the type 17 cytokines, total percentages of IFNγ^+^ cells declined across the subgroups, although there was an increase in IFNγ^+^IL-17A^+^ cells. Taken together, these data support the value of CCR6 as a marker of activity of the broad type 17 program in which type 17 genes show varying degrees and pathways of coordinated vs. independent regulation, demonstrate increasing type 17 and decreasing type 1 and type 2 character across the CCR6-defined memory subgroups, and suggest that the patterns of gene differentiation and resulting phenotypes for the type 17 cells progress along an extended continuum.

**Figure 2.**
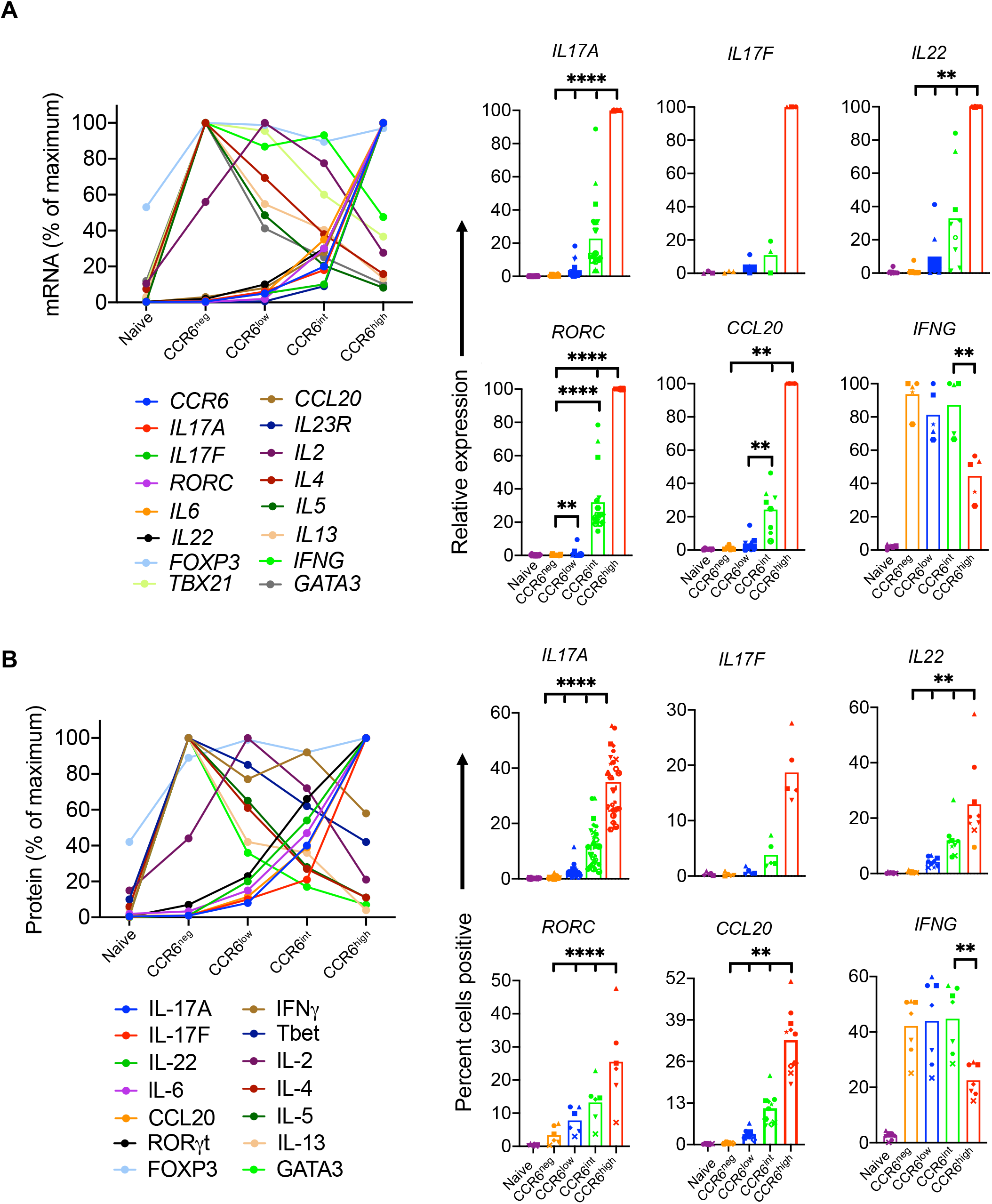
Gradients of expression of lineage-associated genes with increasing CCR6. **(A)** Relative mRNA expression was measured using real-time reverse-transcriptase (RT)-PCR. Ct values of all genes were normalized to *GAPDH*, and relative mRNA expression is shown for means (left panel) or for individual values (right panel) from four or more donors as percentages of the value in the highest expressing subgroup. **(B)** Protein levels in naïve and CCR6-defined memory subgroups for IL-2, IL-4, IL-5 and IL-13 were measured by ELISA from cell supernatants after 12 h of activation with PMA and ionomycin. For other proteins, cells were stimulated with the leukocyte activation cocktail for 6 h, fixed and permeabilized, stained for indicated genes, and analyzed by flow cytometry for percent of cells staining positive. In the left panel, means are shown of values from three or more donors, calculated as percentages of the value in the highest expressing subgroup. The right panel shows percentages of positive-staining cells from five or more donors. Symbols in the right panels indicate data from individual donors and bars indicate means. *p* values were determined using the Wilcoxon matched pairs signed rank test. ^*^*p* < 0.05, \*\**p* < 0.01, ^****^*p* < 0.001.

### The gradient across the CCR6-defined memory subgroups reflects a continuous range of phenotypes of individual cells

For genes/proteins other than CCR6, the patterns that we described for type 17 genes across the CCR6-defined memory subgroups could represent mixtures of homogeneous subsets of, for example, negative and positive cells in varying proportions vs. individual cells with ranges of phenotypes – or some combination thereof. We addressed this issue by using Fluidigm Dynamic Arrays for RT-PCR, which allowed quantitative analysis of expression of a panel of lineage-associated, activation-associated, and housekeeping genes in single cells from each of the naïve, CCR6^neg^, CCR6^low^, CCR6^int^, and CCR6^high^ memory subgroups. Hierarchical clustering of the normalized expression profiles shows co-expression in single cells of type 1 (*IFNG, TBX21, TNF, CXCR3*; highlighted in blue), T_reg_-associated (*IKZF2* and *FOXP3*; highlighted in green), type 2 (*ETS1, GATA3, IL13, IL4*; highlighted in orange), and type 17 (*RORA, KLRB1, CCR6, IL22, IL17A, IL23R, RORC*, and *CCL20*; highlighted in red) genes (Fig. 3 A). Using mean values of gene expression, calculations of correlation coefficients between pairs of the CCR6-defined memory subgroups revealed the ordered relationship of CCR6^low^ to CCR6^int^ to CCR6^high^ (Fig. 3 B), consistent with the analyses from the bulk subgroups.

**Figure 3.**
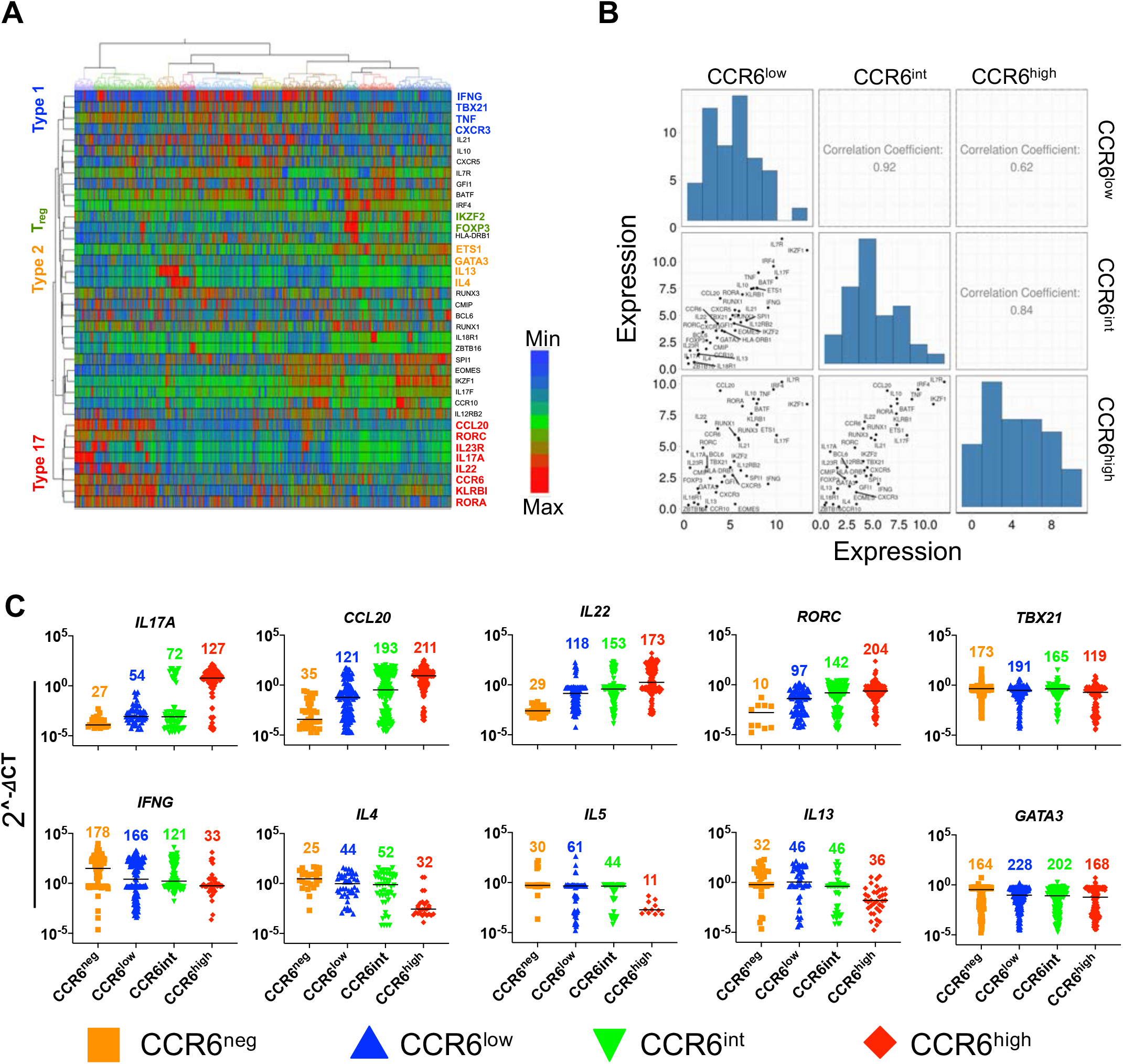
Single-cell data reveal a continuum of expression levels among lineage-associated genes. **(A)** Heat map representation of gene expression profiles using hierarchical clustering showing co-expression of lineage-specific genes in individual cells across naïve cells and CCR6-defined memory subgroups for 282 cells per subgroup. Delta Ct values were calculated using the average of housekeeping genes *GAPDH* and *PGK1*, and after excluding genes with low and/or inconsistent expression, a total of 38 non-housekeeping genes were included in the final analysis. Rows are genes and columns are cells. Some clustered, lineage-associated genes are indicated. **(B)** Correlations among CCR6^low^, CCR6^int^ and CCR6^high^ memory subgroups based on means of expression of the genes in A. **(C)** Gene expression values normalized as described in Materials and Methods for BioMark HD data for the 1410 cells from naïve and CCR6-defined memory subgroups. Each point represents a single cell, and each cell is colored based on its subgroup as indicated. The 1410 cells analyzed in A-C were from three donors.

For the type 17 genes, *IL17A, CCL20, IL22* and *RORC*, the numbers of cells with detectable expression and the levels of expression within single cells increased from the CCR6^low^ to CCR6^high^ subgroups (Fig. 3 C). The data suggest that for the type 1 and type 2 genes, the decreases in expression along the CCR6 continuum in analysis of the bulk population were due predominantly to decreases in the numbers of expressing cells (*IFNG* and *TBX21*), the expression levels per cell (*IL4* and *IL13*), or both (*IL5*). Together these data demonstrate that the gradient of type 17 character among the CCR6-defined memory subgroups at the level of populations reflects a continuum at the level of individual cells, with single cells showing many intermediate phenotypes characterized by a wide range in their expression of signature type 17 genes as well as signature genes of alternative lineages.

### Epigenomes are stable through homeostatic proliferation across the type 17 continuum

We investigated the stability of phenotypes across the type 17 continuum by analyzing genome wide CpG methylation. Methylation of cytosines at CpG dinucleotides is the epigenetic change best associated with stability in patterns of gene expression and reflects the reshaping of these patterns in memory T cells (59, 60). The global methylation profiles reveal a progressive shift from higher to lower CpG methylation from naïve to CCR6^high^ cells (Fig. 4 A). The corresponding unsupervised principal-component analysis shows the large distance between the naïve and memory cells along principal component 1 and the separation along principal component 2 in the order from CCR6^neg^ to CCR6^low^ to CCR6^int^ to CCR6^high^ (Fig. 4 B). Volcano plots of changes in overall methylation levels at individual sites for each of the CCR6-expressing subgroups versus the CCR6^neg^ cells show the predominance of demethylated versus hypermethylated sites from the CCR6^low^ to CCR6^high^ cells and the prominent demethylation at multiple type 17 genes, such as *CCR6, IL-17A, IL-17RE, IL-26, IL-22, CCL20, RORC, RORA, FURIN, SGK1*, and *RUNX3* (Fig. 4 C). To visualize better the changes in methylation status at genes of interest across the CCR6 subgroups, we identified the gene-associated CpG sites showing the greatest variabilities in their beta values across the subgroups and displayed these values in each subset (Fig. 4 D). We found progressively lower overall beta values at sites within multiple type 17 genes, and either little change or progressively higher beta values at sites within type 1 and type 2 genes, such as *CXCR3, IFNG, TBX21, IL4* and *IL13* along the continuum from naïve to CCR6^high^ cells.

**Figure 4.**
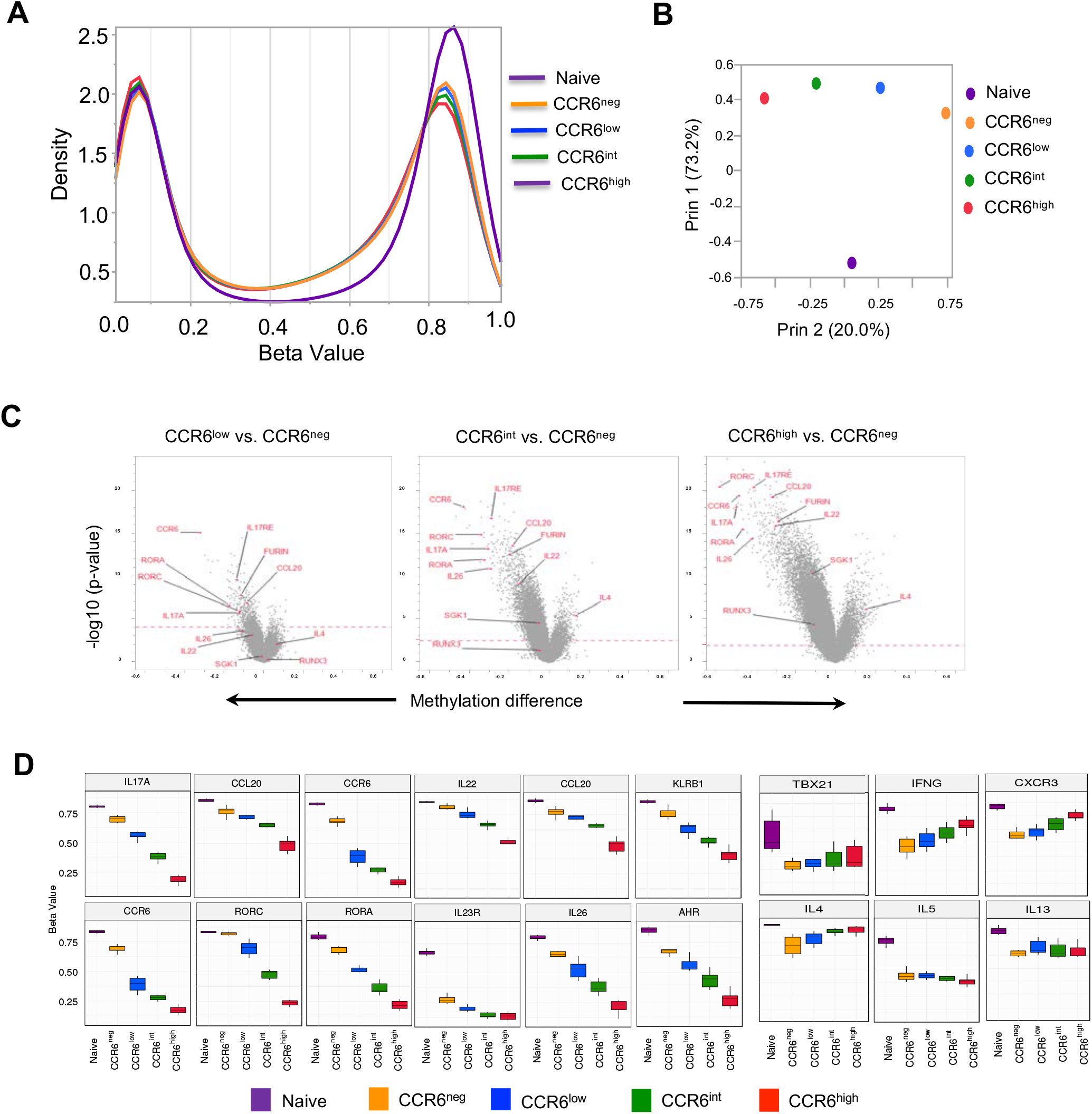
Progressive changes in CpG methylation across CCR6-defined subgroups. **(A)**Distribution of beta values across 450,000 CpG sites using the Illumina Infinium Methylation Assay with naïve cells and the CCR6-defined memory subgroups. Beta values (ratios) of 0.0 and 1.0 correspond to 0 and 100% methylation, respectively, of the sequences at a given CpG site. Data were combined from six donors. **(B)** Correlation and principal variance component projections of naïve cells and CCR6-defined memory subgroups-based genome-wide CpG methylation using the Illumina Infinium Methylation Assay. **(C)** Volcano plots showing differences in methylation beta values at CpG sites in the CCR6^low^, CCR6^int^ and CCR6^high^ subgroups compared with the CCR6^neg^ subgroup versus significance (-log10 p-values). The red dotted line marks the cutoff for significance of differences between beta values at a false discovery rate ≤ 0.05. Some CpG sites at genes of interest are labelled. **(D)** For each of the indicated genes, box plots of methylation beta values are displayed for the single CpG site with the greatest variability among the cell subgroups. For A-D, data from six donors were combined for the analysis.

To challenge the stability of the cells’ phenotypes under homeostatic-like conditions, we activated CFSE-loaded naïve, CCR6^neg^, CCR6^low^, CCR6^int^, and CCR6^high^ cells using low concentrations of anti-CD3 plus homeostatic cytokines for seven days and purified non-proliferated and proliferated cells (Fig. 5 A). To check if there had been a change in the cells’ type 17 character, we compared expression of *IL17A, RORC*, and *CCL20* from day 0 and day 7 proliferated cells after activation with PMA and ionomycin. Proliferation under these conditions did not change the expression of these genes (Fig. 5 B). We analyzed methylation states in cells at day 0 and on day 7 in cells that had either failed to divide (d7NP) or divided three or more times (d7P). In general, principal component analysis positioned the memory cell samples according to their CCR6-defined subgroup in a pattern like that in Figure 4 B, regardless of whether the cells had proliferated (Fig. 5 C). Comparing each of the CCR6-expressing subgroups versus the CCR6^neg^ cells at day 0 (Fig. 5 D, upper panel) showed a similar pattern as in figure 4 C, whereas the methylation levels at individual sites for day 0 versus day 7 proliferated cells for each of the CCR6-expressing subgroups showed no significant changes across the genome (Fig. 5 D, lower panel). Similarly, for genes of interest, methylation at the single CpG sites that displayed the greatest variability across the naïve and CCR6-defined memory subgroups showed no significant changes resulting from culturing or proliferation over 7 days (Fig. 5 E). Together, these data demonstrate that the progressive pattern in lineage-related gene expression across the CCR6-expressing memory subgroups correspond to epigenetic changes at these loci and the genome-wide and gene-specific patterns are unchanged across cell divisions under homeostatic-like conditions, suggesting that the cells’ phenotypes will remain stable during homeostasis *in vivo*.

**Figure 5.**
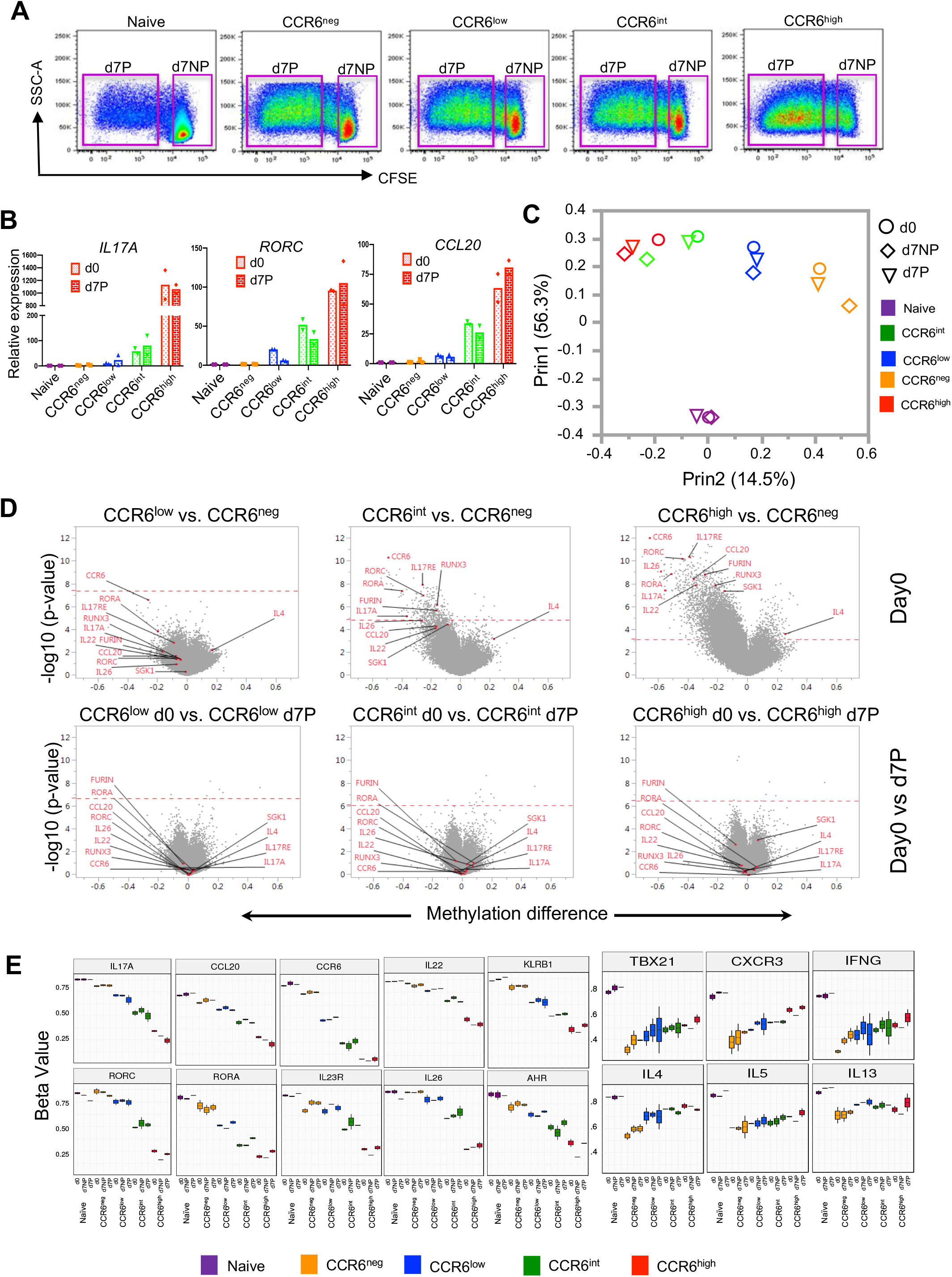
Stable epigenomes of CCR6-defined memory subgroups across cell divisions. **(A)** Distribution of beta values across 450,000 CpG sites using the Illumina Infinium Methylation Assay with naïve cells and the CCR6-defined memory subgroups. Beta values (ratios) of 0.0 and 1.0 correspond to 0 and 100% methylation, respectively, of the sequences at a given CpG site. Data were combined from six donors. **(B)** Correlation and principal variance component projections based on the CpG methylation data of cell subgroups at day 0 (d0) and for cells either non-proliferated (d7NP) or showing three or more cell divisions (d7P) at day 7. **(C)** Volcano plots showing differences in methylation beta values between CpG sites in the CCR6^low^, CCR6^int^ and CCR6^high^ subgroups compared with the CCR6^neg^ subgroup at day 0 (top) or for each subgroup between cells at day 0 and proliferated cells at day 7 (bottom). The red dotted line marks the cutoff for CpG sites with significant differences at a false discovery rate ≤ 0.05. Some CpG sites at genes of interest are labelled. **(D)** Box plots of methylation beta values for CpG sites at lineage-associated genes across cell subgroups at day 0 and for cells either non-proliferated or proliferated at day 7. Data for B-D were combined from two donors.

### CCR6-expressing subgroups share T cell clonotypes with changing frequencies

To investigate the clonal relationships among the CCR6-expressing memory cells we determined the sequences of TCRβ chain mRNAs from naïve cells and CCR6-defined memory subgroups. Using the Morisita-Horn method to compute similarity, the naïve cells show little similarity with the memory cells and the memory subgroups are ordered from CCR6^neg^ to CCR6^low^ to CCR6^int^ to CCR6^high^ (Fig. 6 A, left panels). Circos plots of the 20 most abundant clonotypes in each subgroup reveals that the frequencies of shared clonotypes are most similar between adjacent subgroups and differ most between non-adjacent subgroups (Fig. 6 A, right panels). For example, several clonotypes are found at high frequencies in both the CCR6^int^ and CCR6^high^ subgroups, whereas clonotypes shared between the CCR6^low^ and CCR6^high^ cells that are present at high frequencies in the former are generally at low frequencies in the latter and vice versa.

**Figure 6.**
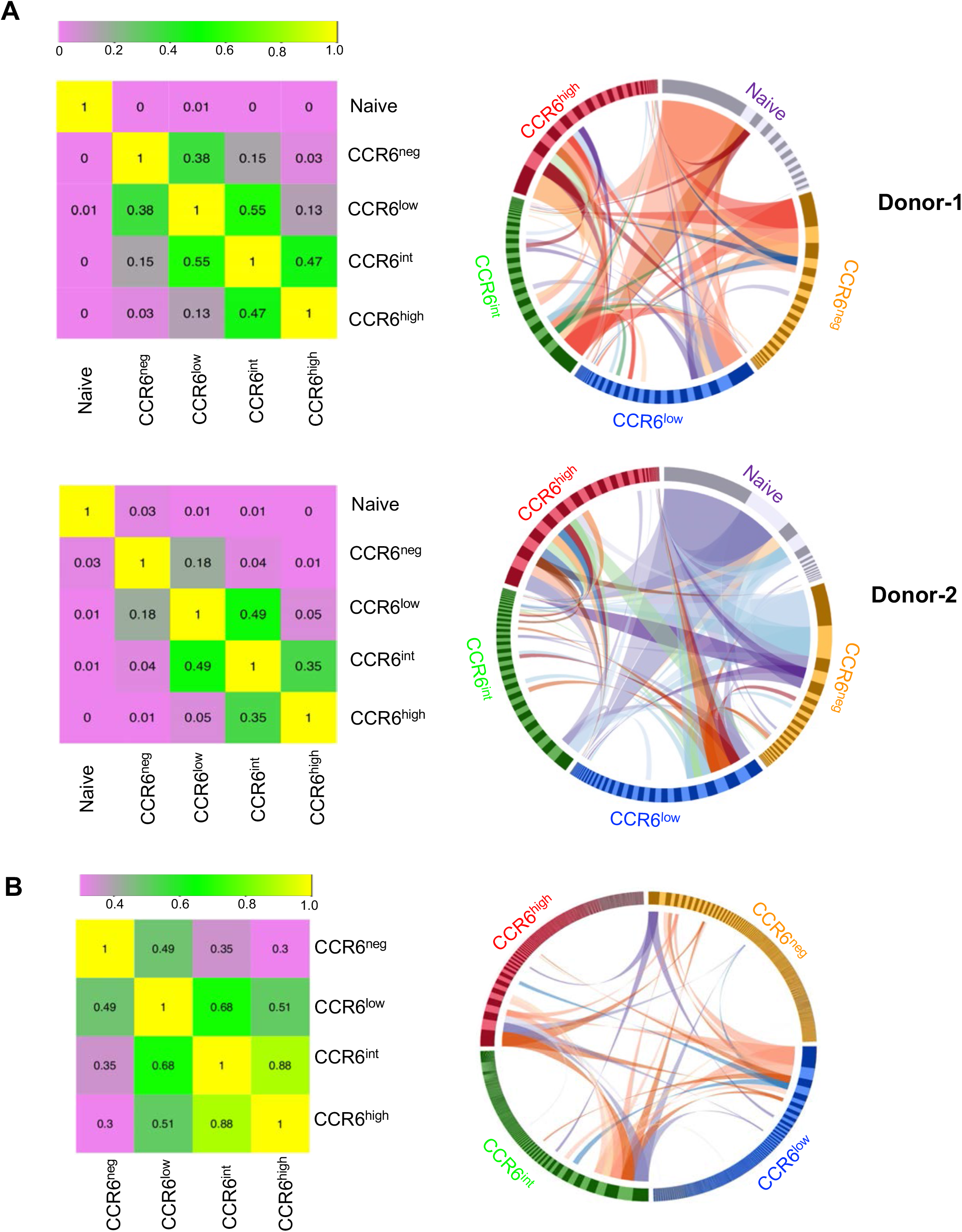
TCR sequencing reveals shared clonotypes whose frequencies change across CCR6-defined memory subgroups. **(A)** Two million T cells from each subgroup from two donors were used for sequencing TCR β chain cDNAs in bulk at a depth of 6.3-9.3 reads per cell and V(D)J sequences were used for identifying clonotypes. Morisita-Horn Similarity Indexes were calculated for pairwise comparisons to yield values between 0 (lowest similarity) and 1 (left panels). Circos plots display all clonotypes with the twenty highest frequencies in any subgroup (right panels). Bars in the circle have widths that are proportional to clonotype frequencies and are arranged within the cell subgroups clockwise in order of decreasing frequencies. Ribbons show connections between clonotypes among the cell subgroups for clonotypes with frequencies of 0.04 or greater within any cell subgroup. **(B)** Cells from each CCR6-defined memory T cell subgroup from a CMV-seropositive donor that proliferated in response to CMV antigens were used for single-cell sequencing of TCR α and β chains. Paired productive sequences were obtained from approximately 4000-6000 cells from each subgroup. Morisita-Horn Similarity Indexes were calculated for pairwise comparisons as in A (left panel). A Circos plot was produced using all clonotypes (right panel). Bars in the circle have widths that are proportional to clonotype frequencies and are arranged within the cell subgroups clockwise in order of decreasing frequencies. Ribbons show connections between clonotypes among the cell subgroups for clonotypes with frequencies of 0.04 or greater within any cell subgroup.

In addition to analyzing clonotypes using TCRβ chain sequences from the bulk populations, we determined clonotype identities with increased specificity by sequencing both TCRα and TCRβ chain mRNAs from single cells. Given the limitations in sample sizes with this method and the significant diversity of the repertoire of non-selected memory T cells, we analyzed cells from the CCR6-defined memory subgroups purified from a donor seropositive for CMV based on the cells’ proliferative responses to CMV antigens (Fig. S3 A). We processed a total of 27,645 cells and recovered paired, productive sequences for TCRα and TCRβ chains from approximately 67% to 71% of the cells, depending on the subgroup (Fig. S3, B and C), and only these sequences were used for subsequent analysis. Computing similarities again ordered the subgroups from CCR6^neg^ to CCR6^low^ to CCR6^int^ to CCR6^high^ (Fig. 6 B, left panel). Circos plots including all the clonotypes (Fig. 6 B, right panel) or the 20 most abundant clonotypes (Fig. S3 D) show that frequencies of shared clonotypes are most similar between adjacent subgroups and differ most between non-adjacent subgroups. These analyses of both the non-selected and CMV-reactive clonotypes reinforce the validity of considering all the CCR6-expressing Th cells as one family, since they show, for example, that the CCR6^low^ subgroup is closer to the CCR6^int^ subgroup than to the CCR6^neg^ cells and demonstrate the same relationships among subgroups as reflected in the gene and protein expression data. Together, the results suggest that at least some cells within the subgroups originate from common T cell clones whose progeny have differentiated, with dynamic changes in their frequencies, along the continuum of type 17 character. To lend further support the directionality of this pathway we performed single-cell droplet-based mRNA sequencing on cells from the CCR6-expressing subgroups from an additional donor. Trajectory analysis of these cells suggested progression from the CCR6^low^ and CCR6^int^ to the CCR6^high^ cells in pseudotime (Fig. S3 E).

### Correlations among expression levels of lineage-associated genes in single cells

To understand the relationships among the Th lineages across the type 17 continuum we analyzed co-expression of lineage-associated genes in the single cells from the naïve and CCR6-defined memory subgroups. We found that co-expression of type 17, type 1 and type 2 genes is common. From the 251 out of the total 1410 naïve and memory CD4^+^ T cells from three donors where, following activation, we could detect expression of *IL17A*, 44% co-expressed *IFNG*, 33% co-expressed a type 2 cytokine gene (*IL4* and/or *IL5* and/or *IL13*), and 54% co-expressed *IFNG* and/or a type 2 cytokine gene. In analyzing those cells that co-expressed pairs of lineage-associated genes, we compared normalized values of expression (Fig. 7 A). Cells co-expressing type 17 genes generally showed positive correlations in their levels, with cells expressing the highest levels enriched in the CCR6^high^ subgroup. Positive correlations were also found for genes within the type 1 and type 2 gene sets. However, positive correlations were not found for *IL17A* versus type 1 and type 2 genes, and these co-expressing cells were found less commonly in the CCR6^high^ subgroup. For the cells co-expressing *IL17A* and *IL4* and/or *IL13*, virtually no cells co-expressed *IL17A* and *IL4* or *IL13* at high levels.

**Figure 7.**
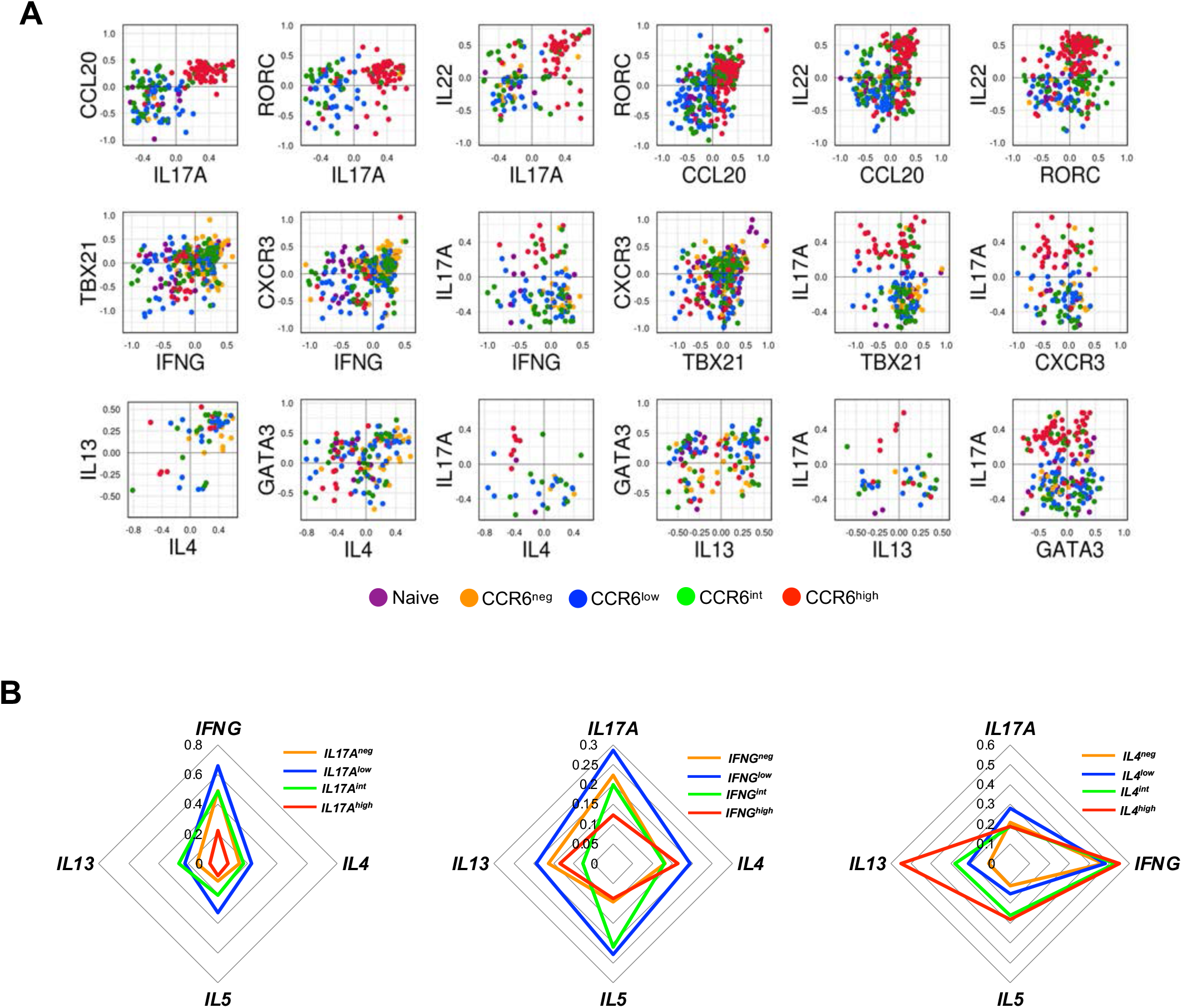
Co-expression of Th lineage-associated genes in single cells. **(A)** Normalized values for expression of the indicated genes from 1410 cells from naïve and CCR6-defined memory subgroups from three donors were pooled and the values from cells co-expressing pairs of genes were plotted for comparison. Each dot corresponds to a single cell and dots are colored based on their subgroups. **(B)** The normalized gene expression data from 1062 cells from the CCR6^neg^, CCR6^low^, CCR6^int^, and CCR6^high^ memory cells from three donors were pooled and cells were divided into positive and negative for *IL17A, IFNG*, or *IL4*, and the positive cells were equally divided into low, intermediate, and high groups based on their expression values for the respective genes. Within each of the four *IL17A*-, *IFNG*-, or *IL4*-defined and color-coded groups of cells, frequencies of cells expressing the other indicated genes are displayed on the concentric diamonds.

In an alternative analysis we combined all the memory cells regardless of their CCR6 expression, and for each of the genes, *IL17A, IFNG* and *IL4* separated cells into four groups, including a group without any expression and three equal-sized groups of low, intermediate, and high expressers. For each group we determined the frequencies of cells with detectable expression of other signature genes. For non-expressing cells or cells that were expressing various levels of *IL17A* the frequencies of cells co-expressing *IFNG* diminished from *IL17A*^low^ to *IL17A*^int^ to *IL17A*^high^ cells (Fig. 7 B), consistent with the trend found in our other data. Surprisingly, however, the *IL17A*^low^ cells had higher frequencies of cells expressing *IFNG*, as well as *IL4, IL5*, and *IL13*, than did the *IL17A*^neg^ cells. For *IL4, IL5*, and *IL13*, the *IL17A*^int^ cells also had higher frequencies of expressers as compared with the *IL17A*^neg^ cells. Patterns that were generally analogous were also found in analyses of cells separated based on their expression of *IFNG* or *IL4*. These data suggest that initial stimulation of CD4^+^ T cells *in vivo* leads to pan-activation of multiple effector pathways, and that these cells can persist in the resting memory population.

### An increase in RORγt can drive type 17 progression from the CCR6^low^ subgroup

The gradients of phenotypes, clonal relationships, and patterns of lineage-gene co-expression that we identified among the CCR6-expressing CD4^+^ T cells suggested that cells could move down this continuum to acquire more type 17 character while diminishing their type 1 and type 2 features. To demonstrate this and understand the outcomes for individual cells we over-expressed *RORC* in CCR6^low^ CD4^+^ T cells by transduction using lentiviral vectors encoding RORγt and GFP or GFP alone and purified GFP-positive cells. The bulk cultures of transduced cells showed increases in mRNAs for *RORC, IL17A* and *CCL20*, a modest decrease in the mRNA for *IFNG*, and more significant decreases for the mRNA for *IL4* (Fig. S4 A).

For determining the effects of transduction of *RORC* on individual CCR6^low^ CD4^+^ T cells we did single cell analysis using Fluidigm Dynamic Arrays for RT-PCR. Comparative data on the frequencies of cells expressing detectable expression of signature genes in *RORC*-virus transduced versus control-virus transduced cells revealed significant increases in *RORC* and *IL17A*, decreases in *IL4, IL5*, and *IL13*, and little change in *IFNG* (Fig. 8 A, left). In analyzing only the cells expressing *IL17A*, the *RORC*-transduced cells showed an increase in frequencies of cells co-expressing *IFNG* and decreases in cells co-expressing *IL4, IL5*, and *IL13* (Fig. 8 A, right). For the *RORC*-transduced cells, violin plots show increases in the frequencies and levels of expression of the type 17 genes *IL17A, IL22, CCL20* and *IL23R* and the opposite effects for *IL4* and *IL13* (Fig. 8 B). We also investigated the “dose response” of cytokine genes to RORγt by comparing the levels of mRNAs of *RORC* to those of the cytokine genes in the *RORC*-transduced cells. The levels of expression of RORC showed positive correlations with expression of *IL17A* and *IL22*. By contrast, even though transduction with *RORC* decreased the frequencies of cells with detectable expression of *IL4*, there was no correlation between the levels of *RORC* and *IL4* mRNAs in individual cells in which expression of *IL4* remains detectable. Expression of *RORC* and *IFNG* also failed to show any correlation (Fig. 8 C). To address whether forced expression of RORγt might be altering frequencies of cells with relevant patterns of gene expression by selective effects on cell proliferation during the 3-day cultures, we analyzed mRNA levels of cytokine genes in transduced and DDAO-loaded cells and found that patterns of cytokine gene expression did not differ as a function of cell divisions, suggesting that RORγt was having direct effects the cells’ phenotypes (Fig. S4 B). Together these data suggest that within the milieu of the CCR6^low^ cells RORγt alone is capable of driving expression of signature type 17 genes, and that increasing expression of RORγt is a mechanism for driving these cells further down the type 17 continuum.

**Figure 8.**
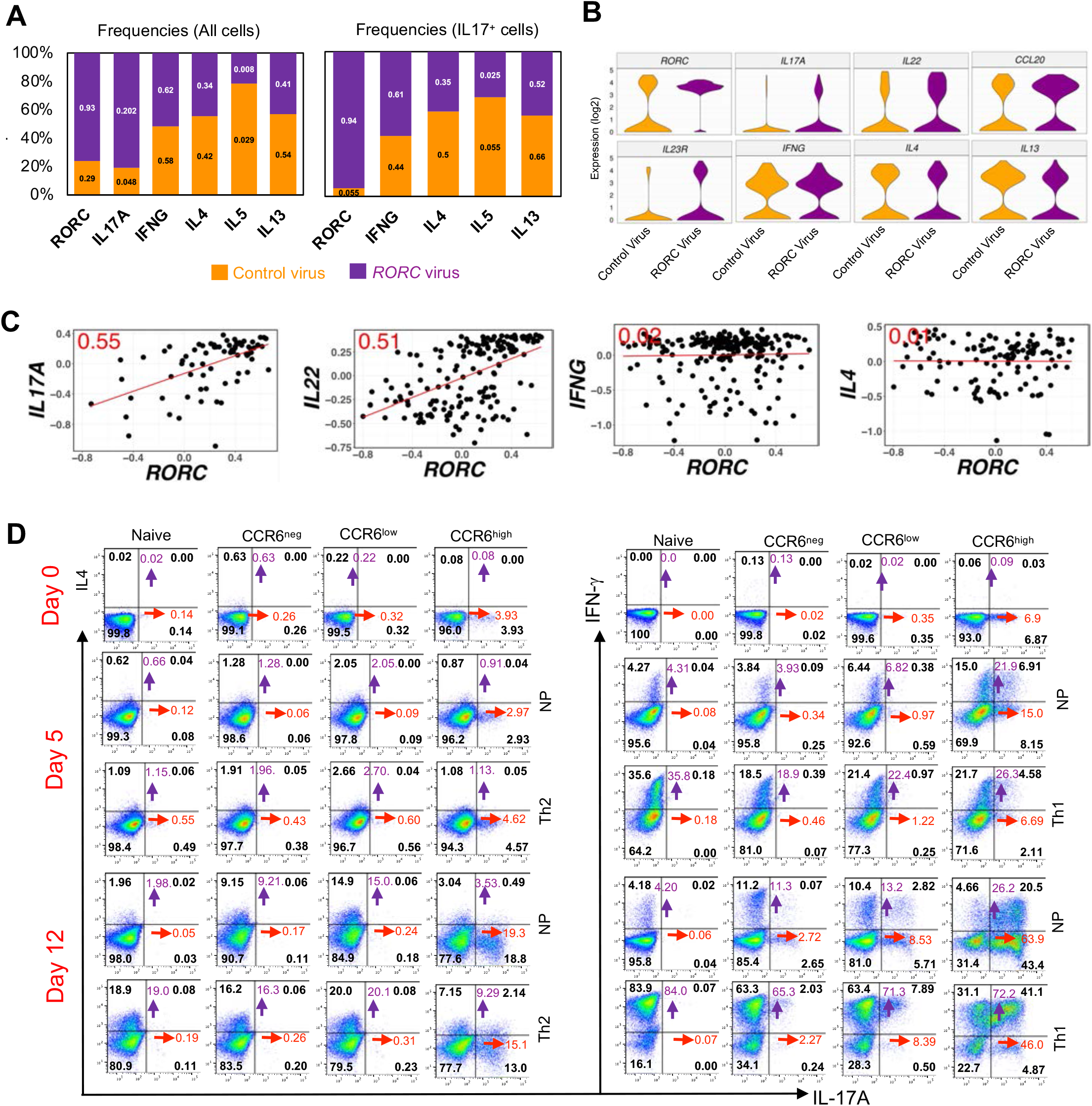
Position in the type 17 continuum determines sensitivity to master regulators and mechanisms of plasticity. **(A)** Numbers on the lower and upper bar segments show frequencies of cells expressing the indicated genes in the control- and *RORC*-transduced cells, respectively. Each bar is divided according to the relative frequencies of the control- and *RORC*-transduced cells expressing the indicated genes. Left panel shows data from all transduced cells and right panel shows data only from cells with detectable expression of *IL17A*. **(B)** Violin plots with expression distributions for the indicated genes in control-or *RORC*-transduced cells. **(C)** Normalized Ct values are plotted for *RORC*-transduced cells that showed co-expression of *RORC* with *IL17A, IL22, IFNG* or *IL4*. Each dot represents a single cell, the red line is the least squares regression line, and the number in red in each plot is the correlation coefficient, *r*. For A-C, data are pooled from four donors. **(D)** Dot plots showing intracellular expression on days 0, 5 and 12 after culturing in non-polarizing (NP), Th2 or Th1 conditions of IL-17A vs IL-4 and IL-17A vs IFN in naïve cells (CD4^+^CD45RO^−^CD25^−^CXCR5^−^CCR4^−^CXCR3^−^), and IL-17A vs IL-4 in CCR6^neg^(CD4^+^CD45RO^+^CD25^−^CXCR5^−^CCR4^−^CCR6^−^), CCR6^low^(CD4^+^CD45RO^+^CD25^−^CXCR5^-^ CCR4^−^CCR6^low^), and CCR6^high^(CD4^+^CD45RO^+^CD25^−^CXCR5^−^CCR4^−^CCR6^high^) subgroups and IL-17A vs IFN in CCR6^neg^(CD4^+^CD45RO^+^CD25^−^CXCR5^−^CXCR3^−^CCR6^−^), CCR6^low^(CD4^+^CD45RO^+^CD25^−^CXCR5^−^CXCR3^−^CCR6^low^), and CCR6^high^(CD4^+^CD45RO^+^CD25^-^ CXCR5^−^CXCR3^−^CCR6^high^) subgroups. Cells were stimulated with the leukocyte activation cocktail for 6 h, fixed and permeabilized and stained for indicated genes. For determining percentages of cells staining positive, quadrants were drawn based on staining with isotype-matched control antibodies. The numbers in black indicate the percentage of cells in each quadrant, whereas purple and red numbers show the sums of percentages of positive cells in the upper two or rightmost two quadrants, respectively. Data are representative of results from three donors.

Given the plasticity reported for Th17 cells in mice, particularly in their transformation into IL-17^−^ IFNγ^+^ cells (36), we investigated plasticity among the CCR6-defined subgroups by activating and culturing them under non-polarizing, Th2, or Th1 conditions. Starting cells were either CCR4^−^ or CXCR3^−^ to eliminate cells with pre-existing activity of the type 2 and/or type 1 programs, including all cells able to make IL-4 or IFNγ, respectively (61). In all experiments, IL-17A^+^ cells were found principally in the CCR6^high^ subgroup. In the Th2 experiments, there was little difference in frequencies of IL-17A^+^ cells in the Th2 vs. non-polarizing conditions, and IL-17A^+^ cells generally remained IL-4^−^. At 12 days, non-polarizing cultures showed prominent increases in percentages of IL-4-producing cells in the CCR6^neg^ and CCR6^low^ subgroups as compared with the naive and CCR6^high^ cells. The Th2-polarizing conditions resulted in further increases in IL-4^+^ cells, with the fewest IL-4^+^ cells found in the CCR6^high^ subgroup. However, the fold increase in IL-4 production in Th2 vs. non-polarizing conditions was greater in the CCR6^high^ (and naïve) cells as compared with the CCR6^neg^ and CCR6^low^ subgroups (Fig. 8 D, left panels).

In the Th1 experiments, a significant percentage of CCR6^high^ cells, but fewer cells in the CCR6^low^ or other subgroups, acquired the ability to produce IFNγ within five days under non-polarizing conditions. At the same time, both at 5 and at 12 days the fold increases in IFNγ production in Th1 vs. non-polarizing conditions were smaller in the CCR6^high^ cells as compared with the CCR6^neg^ and CCR6^low^ subgroups. Th1 conditions also led to a loss in IL17^+^ IFNγ-cells and, by day 12,a shift of most IFNγ^+^ cells into the IFNγ^+^ IL17^+^ subset (Fig. 8 D, right panels). When considered together, these experiments suggest that progression along the type 17 continuum results in a diminished ability to express the type 2 but not the type 1 program. Of greater interest, the comparisons between cells cultured under non-polarizing vs. polarizing conditions suggest that prior conditioning vs. activating environments contribute differentially to the expression of the programs for the non-Th17 lineages depending on the program and a cell’s position along the type 17 continuum. The cells’ prior imprinting contributes more than subsequent environmental inputs for the CCR6^low^ than CCR6^high^ cells in acquiring an ability to produce IL-4, whereas prior imprinting contributes more than the environmental inputs for the CCR6^high^ than CCR6^low^ cells in acquiring an ability to produce IFNγ.

In an alternative approach to studying plasticity, we transduced CCR6^low^ and CCR6^high^ memory cells (here without removing cells expressing CXCR3 or CCR4) using GFP-expressing vectors encoding *RORC, TBX21*, and *GATA3* and a control vector expressing GFP alone. All experiments used GFP^+^ cells purified by FACS. We checked the activities of these transcription factors initially by transducing CCR6^low^ cells and analyzing mRNAs in the bulk populations. Given the levels of the endogenous mRNAs, the fold increase in the *RORC* mRNA was much greater than for the mRNAs for *TBX21* and *GATA3* (Figure S4 C). As in the experiments described above, transduction with *RORC* led to a large increase in the expression of *IL17A*. Transductions with *TBX21* and *GATA3* led to increased expression of *IFNG* and *IL4*, respectively. We next transduced CCR6^low^ and CCR6^high^ cells with these transcription factors and analyzed mRNAs from individual cells by RT-PCR (Figure S4 D). For *IL17A*, transduction with *RORC* led to large increases in the frequency of expressing cells and levels in the individual expressing cells in the CCR6^low^ subgroup, and a small increase in the CCR6^high^ subgroup. For each of the mRNAs, *IFNG, GATA3*, and *IL4*, transduction with *RORC* led to a decrease in the CCR6^low^ subgroup and little change in the CCR6^high^ subgroup, with the possible exception of some decrease in *IL4* expression also in the latter cells. Transduction with *TBX21* led to modest increases in the frequency and levels (in the individual expressing cells) of *IFNG* in the CCR6^low^ subgroup, but no changes in CCR6^high^ cells, and no consistent changes in expression of other genes. Transduction with *GATA3* led to modest increases in the frequency and levels of *IL4* in the CCR6^low^ subgroup, but no changes in CCR6^high^ cells, and no consistent changes in expression of other genes. Taken together, these data reveal significant responsiveness for CCR6^low^ cells, but a diminished ability to affect the expression of lineage-specific genes in the CCR6^high^ cells by overexpression of signature transcription factors, suggesting, as we found in our *ex vivo* cross-polarization experiments, that mechanisms of plasticity differ depending on the cells’ positions along the type 17 continuum. For example, the CCR6^low^ cells, which are more responsive to Th1 polarizing conditions, are also more responsive to overexpression of *TBX21*, whereas the CCR6^high^ cells are those showing the greatest prior imprinted ability to upregulate IFNγ while being insensitive to the overexpression of *TBX21*.

## DISCUSSION

Our studies focused on understanding the composition and organization of the memory type 17 Th cell population in healthy humans. One central finding is that these cells, including the progeny of single clones, lie at different positions along an extended continuum of type 17 differentiation on which is superimposed the co-activation of alternative lineages. Although co-expression of alternative lineages diminishes along the continuum, such co-expression is more the rule than the exception and contributes to the population’s extensive heterogeneity. Based on a wealth of data from mouse studies establishing a direct relationship between effector and memory T cells (8, 12-18), we presume that the phenotypes of the cells within this continuum reflect pathways of effector cell activation and differentiation. Our data suggest that these *in vivo* pathways are initiated by activating changes at genes associated with multiple effector lineages, and that the less differentiated type 17 memory cells bearing these changes are particularly capable of expressing a variety of alternative effector phenotypes.

The character of the type 17 continuum was revealed by analyzing cells that differed in their expression of the trafficking/chemokine receptor, CCR6. Studies in mice and humans have suggested that there is a family of Th17-related cells that cannot be identified by relying on the expression of a single cytokine, particularly when limited, as in human studies, to examining cells at one point in time (35, 42). Additionally, the increasing appreciation of memory T cell diversity has led to proposing that the phenotypes of these cells be viewed along multiple vectors, with trafficking patterns representing a major defining feature (62). Whether all CCR6-expressing Th cells belong within a single type 17 family has been questioned (38, 63). However, *in toto*, our data strongly suggest that CCR6 does, in fact, identify a subgroup of related cells, and the expression of *IL17A, IL17F, IL22, CCL20, IL23R* and *RORC* almost exclusively within the CCR6^+^ subset of memory T cells (46-48, 50) and this manuscript), suggests that CCR6 is an appropriate marker for human type 17 cells understood more broadly. We reasoned that analyzing this more inclusive collection of type 17 cells would reveal relationships that might be obscured by studies of the Th17 population narrowly construed, and thereby illuminate both the pathways for differentiation and the nature of inter-lineage relationships in human Th cells *in vivo*.

Analysis of individual cells along the gradient of CCR6 expression showed that expression levels of type 17 genes vary, generally in a coincident fashion, from cell-to-cell, with most type 17 genes showing continua of expression across a wide range of values. High-dimensional studies have identified linear patterns of differentiation with continua of phenotypes as general features of human CD8^+^ (64) and CD4^+^ (65) memory cell populations, and of mouse Th cells activated under polarizing conditions (66).

If, as we presume, the memory cells within the CCR6-expressing subgroups are derived from activated cells that had differentiated to various points along a type 17 continuum, then we might expect the subgroups to share T cell clones in a pattern reflecting the subgroups’ relationships. From both the sequences of β chain mRNAs from the cells analyzed in bulk and the sequences of paired α/β chains of CMV-reactive single T cells we found many shared clonotypes among the memory cell subgroups with changes in frequencies of clonotypes from subgroup to subgroup. The clonotype compositions of the CCR6-negative cells were closest to the CCR6^low^ cells, and the ordered relationship among the CCR6-expressing subgroups matched the pattern derived from the gene and protein expression data, CCR6^low^ to CCR6^int^ to CCR6^high^, consistent with the descendants of individual clones having progressed to different points during the process of activation-driven differentiation.

To investigate the stability of the type 17 continuum, we analyzed CpG methylation in the naïve and CCR6-defined memory subgroups, both in cells immediately after isolation and in cells that had undergone proliferation after gentle activation under non-polarizing conditions *in vitro*. Epigenetic changes have been studied for their roles in establishing and maintaining T-cell memory (67); genome-wide changes in DNA methylation are characteristic of effector/memory differentiation in mouse (68, 69) and human (65) Th cells; demethylation at lineage-specific cytokine genes has been shown to be important for their expression in differentiating Th cells (69, 70); and the preservation of patterns of gene expression in memory T cells has been associated with the stability of gene-specific demethylation across multiple cell divisions *in vitro* and/or *in vivo* (17, 19, 60, 71). Of particular relevance for our studies, demethylation of a region at the *CCR6* promoter was shown to be important for stable expression of CCR6 in human Th cells (59).

We found an overall loss of methylated CpG’s along the CCR6-defined continuum, consistent with increasing memory cell character, and progressive decreases in methylated CpG’s at multiple type 17 genes together with increasing methylation at type 1 and type 2 genes. The patterns of inducibility of type 17 genes and degrees of their site-specific CpG methylation as analyzed at the level of cell populations were also unaltered across the CCR6-expressing memory subgroups following proliferation *in vitro*. These results suggest that the phenotypes of the cells within the CCR6-expressing subgroups are stable under homeostatic conditions as expected for a resting memory population.

The stability of the type 17 continuum under homeostatic conditions notwithstanding, the strong suggestion from the linear relationship among the phenotypes and the clonotype data was that cells could progress along the continuum under the proper conditions of activation. We found that, indeed, a forced increase of expression of RORγt in the CCR6^low^ cells up-regulated the expression of type 17 genes and down-regulated expression of some type 1 and type 2 genes. These changes were evident both in analyzing the cells in bulk and at the level of individual cells, and the single-cell data showed that expression levels of *RORC* mRNA correlated with the levels for a subset of type 17 genes, such as *IL17A* and *IL22*, suggesting that varying expression of *RORC* is one mechanism for establishing the cell-to-cell gradients in gene expression and for moving cells along the continuum.

In our analysis of gene expression levels in single, freshly isolated cells we found positive correlations for genes within the type 17, the type 1, and type 2 lineages, and negative correlations for genes across lineages, especially when a gene from one lineage was expressed at high levels. Overall, cross-suppression was most notable between the type 17 and type 2 pathways, as reflected in the single cell data in which almost no cells co-expressed high levels of both *IL17A* and *IL4* or *IL13*. Nonetheless, co-expression of signature genes of different lineages was the common, with most cells that had detectable expression of *IL17A* co-expressing one or more of the genes that are characteristic of alternative lineages. Although we cannot know if each cell with measurable expression of a given gene could produce the encoded protein in amounts that are biologically relevant, at a minimum, the detection of these mRNAs indicates epigenetic changes at the gene loci that reflect activation of the lineage-specific pathways. For mouse Th cells, complex continua of mixed lineage phenotypes have been described after activation *in vitro* with combinations of “opposing” cytokines (72) and cells with stable mixed lineage phenotypes have been identified from inflammatory reactions *in vivo* (66). Human Th cells co-expressing IL-17A and IFNγ (46-48) have been well described, and co-expression of IL-17A and type 2 cytokines has also been reported (11).

Of particular interest, our analysis of single cells showing different levels of expression of lineage-associated cytokine genes revealed that the frequencies of cells expressing *IFNG* or *IL4* were higher among those cells expressing low levels of *IL17A* than among cells in which expression of *IL17A* was undetectable. Analogous patterns were found for cells with undetectable versus low-level expression of *IFNG* or *IL4* in their expression of the cytokine genes of alternative lineages. These data suggest that initial activation of Th cells *in vivo* leads preferentially to pan-activation of effector pathways and that “early” cells with this history persist within the memory population. Previous experiments analyzing the initial events in the differentiation of activated Th cells have shown, for example, that genes for both Th1 and Th2 transcription factors and cytokines are activated before polarized patterns are established (73). During an acute challenge, when time is of the essence, pathway pan-activation could jump-start effector functions until the response can be focused appropriately. In studies of pathogen-reactive human memory Th cells, the analysis of clonotypes has strongly supported the model of “one cell, many fates”, suggesting that the dominance of a particular Th lineage in response against a pathogen results principally from the differential expansion of a clonotype within a given lineage rather than initial commitment being restricted to a single lineage (11). Within a recall response, such a process would follow naturally from individual memory cells with the phenotypes of pan-activation that we have described. Such cells would maintain within the memory pool the capabilities resulting from the previous multi-pathway co-activation and form cellular substrates with the flexibility to expand along the optimal pathway(s) to meet the demands of host defense.

Co-expression of genes in the type 17 and alternative Th pathways raises the issue of memory/effector Th cell plasticity (74). Soon after the identification of Th17 cells it was reported that they showed an unusual degree of plasticity, particularly in the apparent conversion of mouse IL-17A/F-producing Th cells to cells that were making IFNγ and often no IL-17A/F, and that these “transdifferentiated” cells were mediators of organ injury in models of autoimmune disease (33, 34, 75).

We addressed plasticity in human type 17 cells by activating and culturing naïve and/or CCR6-defined subgroups of memory cells under non-polarizing and polarizing conditions and by transducing these cells with genes for the “master regulators” RORγt, T-bet, and GATA3. In CCR4^−^ memory cells, none of which can produce detectable IL-4 initially, the CCR6^high^ subgroup contained fewer cells producing IL-4 after culturing cells under Th2-polarizing conditions and was less able to increase numbers of *IL4*-expressing cells after transducing with *GATA3* when compared with cells in the CCR6^low^ subgroup. These findings are consistent with the significant progressive decrease in expression of type 2 genes in the freshly isolated cells along the CCR6-defined continuum. However, the contribution of Th2 polarizing conditions on IL-4 production was greater for the CCR6^high^ (and naïve) cells than for the CCR6^neg^ and CCR6^low^ cells, for which much of the ability to produce IL-4 was acquired in the non-polarizing environment. In CXCR3^−^ memory cells, none of which can produce detectable IFNγ initially, a notable finding was the ready acquisition by the CCR6^high^ cells of the ability to produce IFNγ after activation under non-polarizing conditions. For these cells, the added effect of a Th1-polarizing environment on IFNγ production was significantly less marked than it was for the CCR6^neg^ and CCR6^low^ (and naïve) cells.

The results of our polarization experiments suggest two distinguishable processes at work. The first process is “imprinted partial plasticity” as proposed by Crotty, whereby the potential to express aspects of alternative programs have been established during initial activation but are not manifest until re-activation in a permissive environment (38). The second process is plasticity induced by the environmental inputs during reactivation. Our data suggest that the outcomes of reactivating type 17 memory cells will reflect differing contributions of imprinted plasticity and retained flexibility depending on which pathways are activated and, importantly, the cells’ positions along the type 17 continuum.

In summary, we have found that CCR6 identifies a broad but connected family of human type 17 memory Th cells that fall along an extended gradient of type 17 activity. This continuum is evident in cells analyzed in bulk as subgroups or, more importantly, as single cells, and is likely to reflect pathways of clonal differentiation during activation *in vivo*. The epigenomes and phenotypes of cells along the continuum are stable under homeostatic conditions. Nonetheless, increasing expression of *RORC* in CCR6^low^ cells demonstrated a mechanism whereby cells could “move to the right” to acquire additional type 17 character, and cells across the type 17 continuum showed differing mechanisms of plasticity in the inducible expression of signature genes of non-type 17 lineages. Whereas the CCR6^high^ cells were more resistant than the CCR6^low^ cells in activating *IL4*, the CCR6^high^ cells were superior in activating *IFNG* under non-polarizing conditions, representing a novel example of imprinted plasticity that was greater in the more highly differentiated type 17 cells. Considered together, the features of the extended pathway of type 17 differentiation that we have described yield both a linear progression of type 17 activity and a range of other attributes that, importantly, differ as a function of the cells’ positions along the continuum, including mixed-lineage capabilities and varied mechanisms of plasticity to create a multi-functional and adaptable collection of type 17 memory cells. Finally, our sensitive single-cell RT-PCR assays of cytokine gene expression suggest that pan-induction of effector pathways, rather than choice of a single lineage, is a general feature of early Th cell activation *in vivo*, and that cells with the resulting multi-lineage character persist in the memory pool, which may contribute to the memory population’s overall flexibility and adaptability to secondary challenges. These studies provide fundamental information on the composition of the human type 17 memory population and allow for informed inferences and hypotheses as to pathways and mechanisms through which this highly heterogeneous population is formed and can adapt to novel demands of host defense or be misdirected to produce inflammatory damage.

## Acknowledgements

We thank Jinfang Zhu, NIAID, and Jean K. Lim, Icahn School of Medicine at Mount Sinai, for critical reading of the manuscript. We are grateful to Calvin Eigsti and members of the Research Technologies Branch, NIAID, for their help with cell sorting. This research was supported by the Intramural Research Program of the NIH.

