## Supplemental Figures for "Human CCR6^+^ Th cells show both an extended stable gradient of Th17 activity and imprinted plasticity"

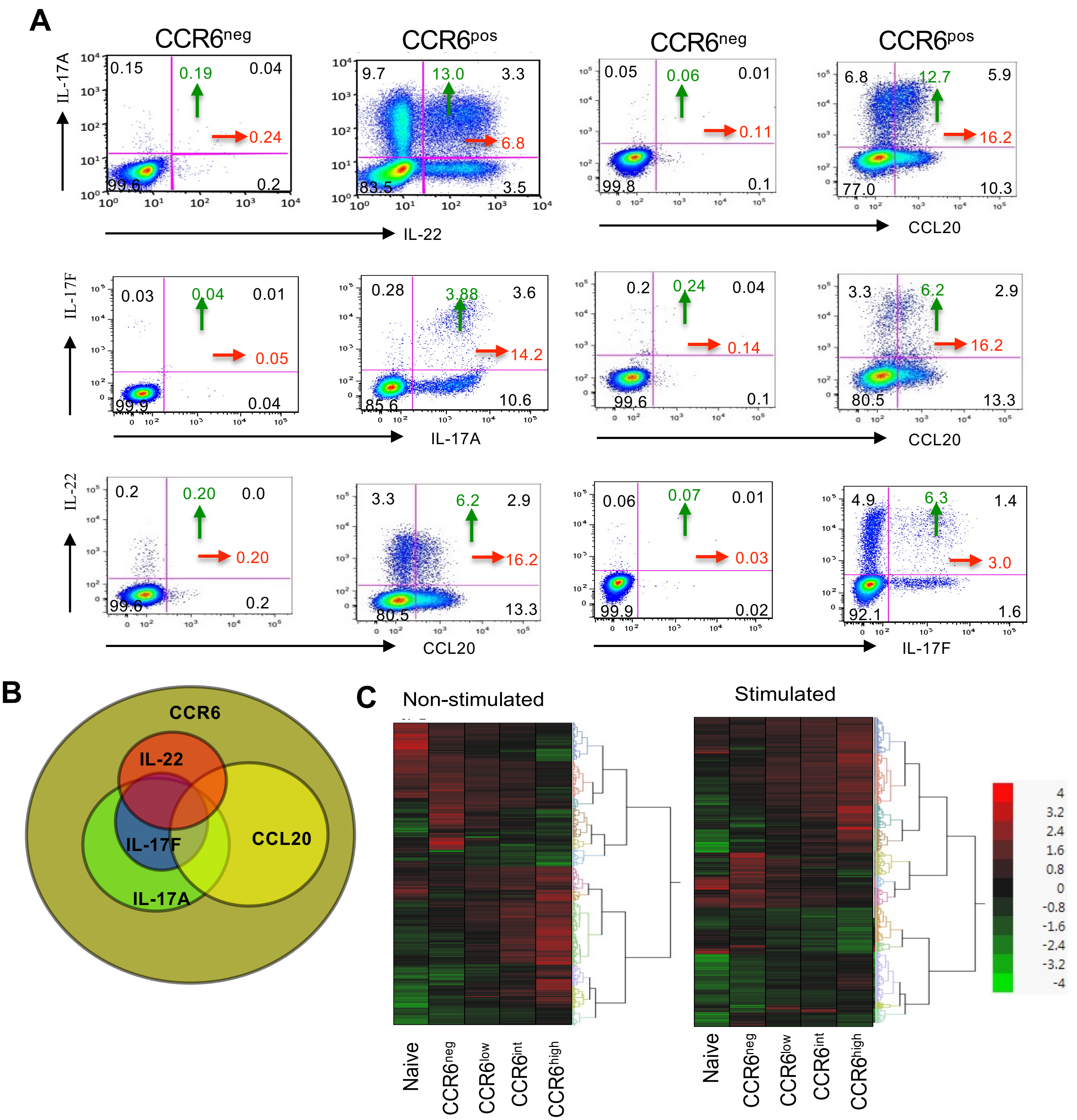


Figure S1. **All Th cells making type 17 cytokines express CCR6.** **(A)** Sorted CCR6$^{neg}$ and CCR6$^{pos}$ cells were stimulated with PMA and ionomycin and stained for the indicated cytokines after permeabilization and fixation. Quadrants were drawn based on staining with isotype-matched control antibodies. The numbers in black indicate the percentages of cells in each quadrant, whereas green and red numbers show the sums of percentages of cells positive in the upper two or rightmost two quadrants, respectively. Results are shown from a single donor, representative of three or more donors. **(B)** Venn diagram shows relationships among cells expressing CCR6 and showing intracellular staining for the indicated cytokines. Results are based on data from three or more donors. **(C)** Expression profiles of differentially expressed genes in CCR6-defined subgroups. The gene-specific probes with *p* values below the false discovery rate in any pairwise comparison among the subgroups are shown in heatmaps. Data in C is combined from eight individual donors.

**Figure S1**

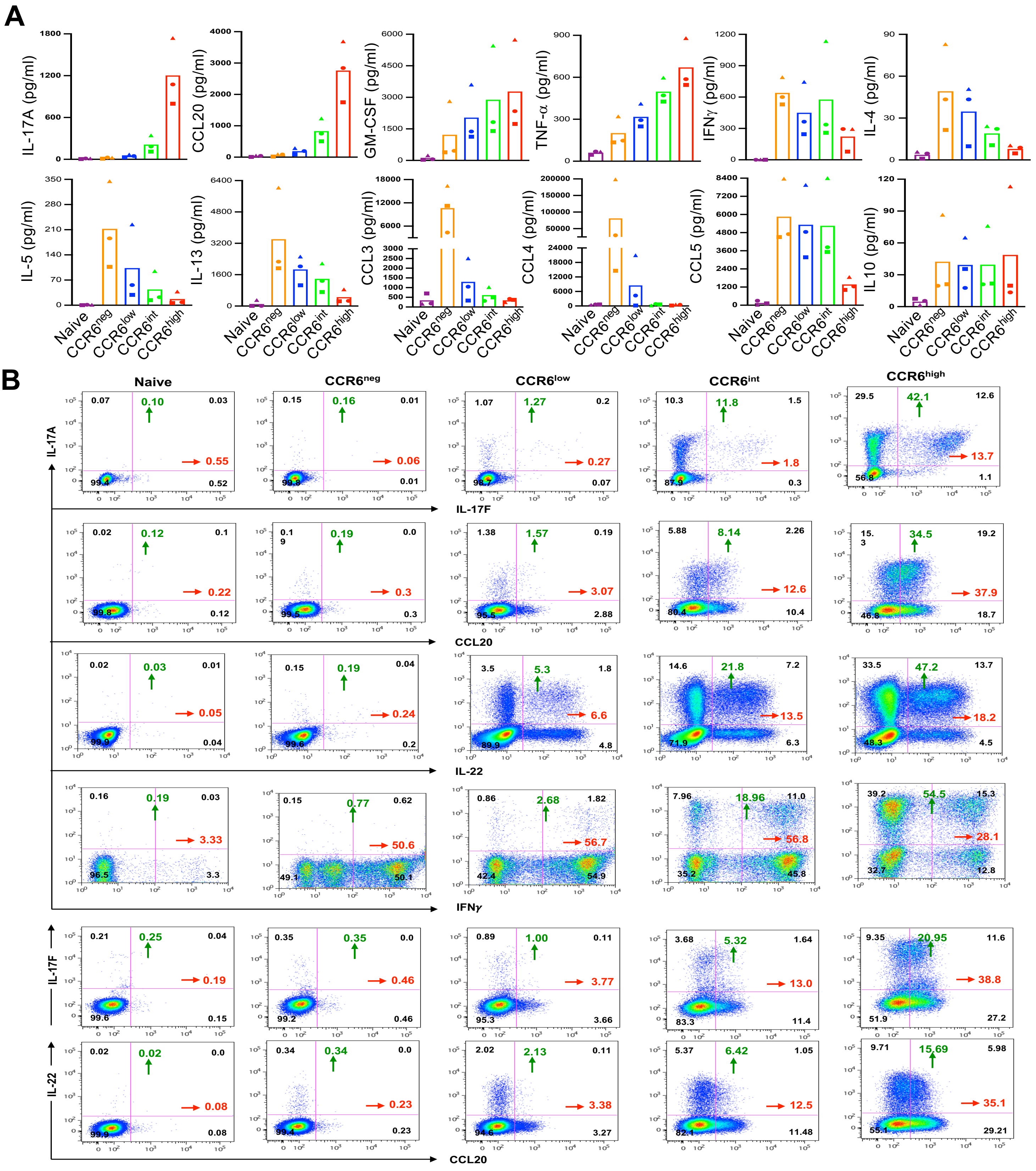


Figure S2. **Cytokine and chemokine production and co-expression by CCR6-defined memory subgroups. (A)** Cytokines and chemokines were measured by ELISA in culture supernatants after cells had been activated for 12 hours with PMA and ionomycin. Bars indicate means. Data are from three donors. **(B)** Density-colored flow cytometry dot plots showing intracellular expression of cytokines and chemokines in FACS-sorted naïve cells and CCR6-defined memory subgroups 6 hours post PMA and ionomycin stimulation. For determining percentages of cells staining positive, quadrants were drawn based on staining with isotype-matched control antibodies. The numbers in black indicate the percentages of cells in each quadrant, whereas green and red numbers show the sums of percentages of positive cells in the upper two or rightmost two quadrants, respectively. Data are representative of results from three or more donors.

**Figure S2**

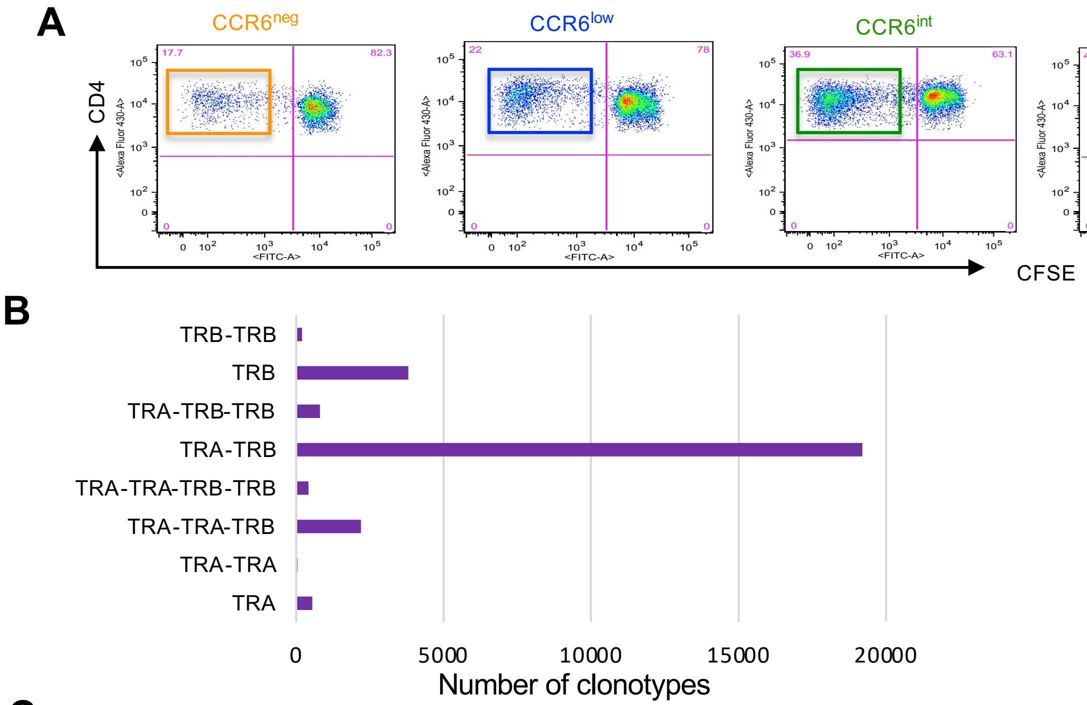


C

| Sample | TCRα (%) | TCRβ (%) | Paired productive chain (%) |
|---|---|---|---|
| CCR6^neg | 4991/5970 (83.6) | 5870/5970 (98.3) | 4117/5970 (68.9) |
| CCR6^low | 6103/8245 (74.0) | 7875/8245 (95.5) | 5546/8245 (67.2) |
| CCR6^int | 6345/7364 (86.2) | 7203/7364 (97.8) | 5219/7364 (70.8) |
| CCR6^high | 5168/6066 (85.2) | 5952/6066 (98.1) | 4318/6066 (71.2) |

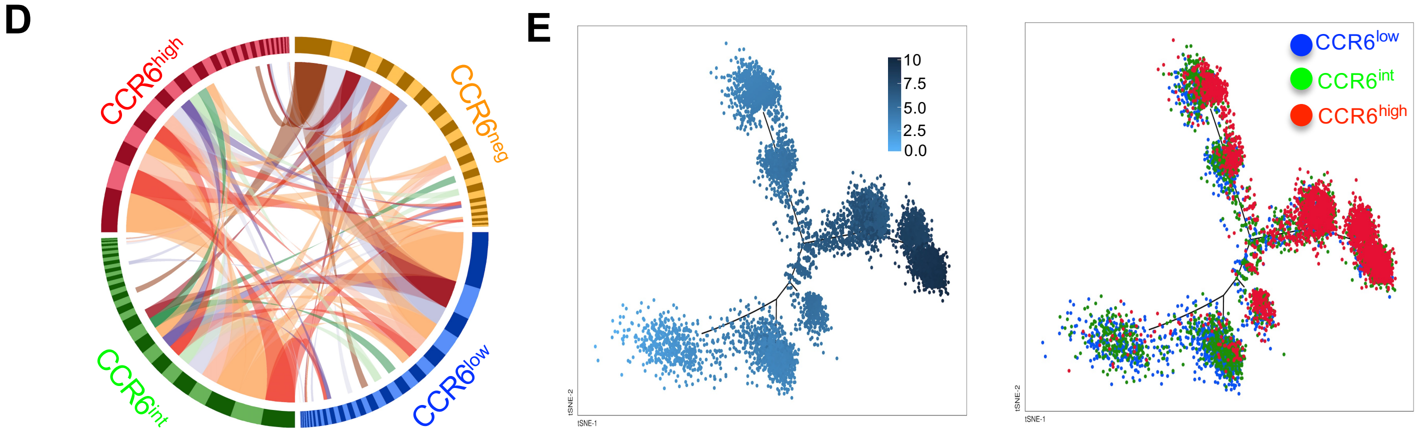


Figure S3. **Shared clonotypes of varying frequencies across the CCR6-defined memory subgroups revealed by analysis of single CMV-reactive cells. (A)** Density-colored flow cytometry dot plots show gating strategy for cell sorting of CMV-reactive, CFSE$^{neg/low}$ cells from CCR6-defined memory subgroups from a seropositive donor. Only CD3$^{+}$CD4$^{+}$ cells are displayed. **(B)** Bars show numbers of clonotypes of each composition among the combined cells from all the cell subgroups. **(C)** Numbers and frequencies of cells with productive TCRa and/or TCRb chains for each of the cell subgroups. **(D)** Circos plot displaying all clonotypes with the twenty highest frequencies in any cell subgroup. Bars in the circle have widths that are proportional to clonotype frequencies and are arranged within the cell subgroups clockwise in order of decreasing frequencies. Ribbons show connections between clonotypes among the cell subgroups for clonotypes with frequencies of 0.04 or greater within any cell subgroup. Data in A-D are from cells from a single donor. **(E)** Cells from the CCR6$^{low}$, CCR6$^{int}$ and CCR6$^{high}$ memory T cell subgroups from one donor were used for single-cell sequencing of mRNAs using a 5' gene expression assay, the data analyzed using Monocle2, and the trajectory displayed using a tSNE plot . Each dot corresponds to a single cell. The left panel coloration reflects pseudotime with 0 assigned to the origin. The right panel shows the positions of cells colored according to CCR6 subgroups.

**Figure S3**

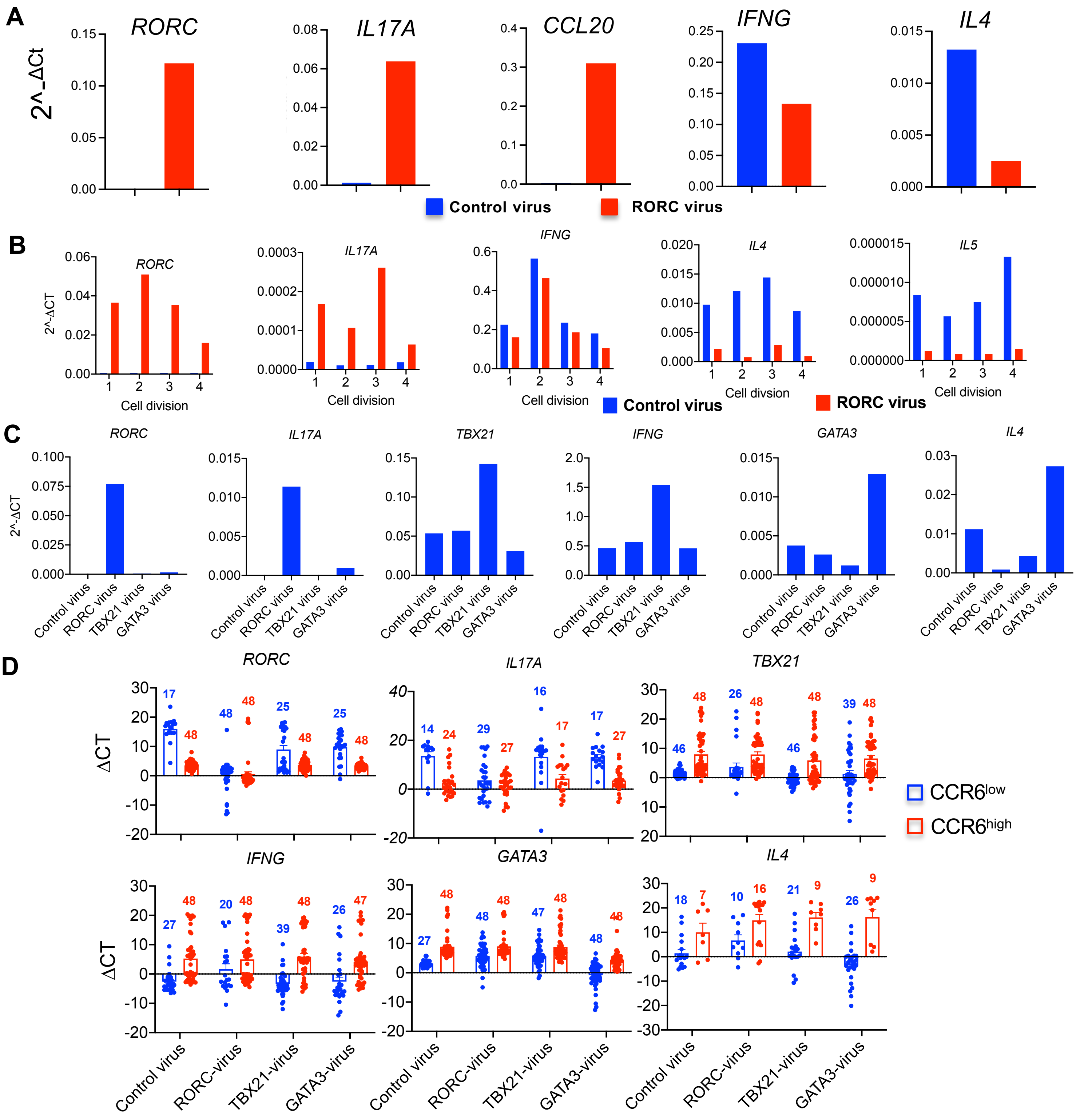


Figure S4. **Plasticity of the CCR6$^{low}$ subgroup revealed by forced expression of *RORC*, *TBX21* and *GATA3*. (A)** Expression of the mRNAs from the indicated genes, measured using RT-PCR, in GFP$^{+}$ control- and *RORC*-transduced cells. Values for these genes were normalized to values for *GAPDH* for calculating delta Ct. Data are from one donor, representative of three. **(B)** CCR6$^{low}$ cells were loaded with DDAO before being used for transduction. After 72 hours, GFP$^{+}$ control- and *RORC*-transduced cells were purified by FACS based on DDAO dilutions. Bars show expression of the mRNAs from the indicated genes, measured using RT-PCR. Values for these genes were normalized to values for *GAPDH* for calculating delta Ct. Data are from one donor, representative of three. **(C)** Expression of the mRNAs from the indicated genes, measured using RT-PCR, in GFP$^{+}$ control-, *RORC*-, *TBX21*-, or *GATA3*-transduced cells. Values for these genes were normalized to values for GAPDH for calculating delta Ct. Data are from one donor, representative of two. **(D)** Expression of the mRNAs from the indicated genes, measured using single-cell RT-PCR, in GFP$^{+}$ control-, *RORC*-, *TBX21*-, or *GATA3*-transduced cells. Values for these genes were normalized to values for GAPDH for calculating delta Ct. Each dot represents a single cell. Bars show means +/- S.E.M. Numbers above each bar indicate the number of cells in that bar with detectable expression of the indicated gene, with maximum number being 48. Data are combined from 24 cells from each of two donors. For B-D, cells were stimulated with PMA and ionomycin for 3 hours.

**Figure S4**
